# Mechanism of forced-copy-choice RNA recombination by enteroviral RNA-dependent RNA polymerases

**DOI:** 10.1101/2025.02.07.637143

**Authors:** Jamie J. Arnold, Alexandre Martinez, Abha Jain, Xinran Liu, Ibrahim M. Moustafa, Craig E. Cameron

## Abstract

Forced-copy-choice recombination occurs at the end of a template, differing from copy-choice recombination, which happens at internal positions. This mechanism may produce full-length genomes from fragments created by host antiviral responses. Previous studies from our laboratory demonstrated that poliovirus (PV) RNA-dependent RNA polymerase (RdRp) switches to an “acceptor” template *in vitro* when initiated on a heteropolymeric RNA-primed “donor” template. Surprisingly, recombinants showed template switching from the 3’-end of the donor template. We have developed a primed-template system to study PV RdRp-catalyzed forced-copy-choice RNA recombination. PV RdRp adds a single, non-templated nucleotide to the 3’-end of a blunt-ended, double-stranded RNA product, forming a “plus-one” intermediate essential for template switching. Non-templated addition of CMP was favored over AMP and GMP (80:20:1); UMP addition was negligible. A single basepair between the plus-one intermediate and the 3’-end of the acceptor template was necessary and sufficient for template switching, which could occur without RdRp dissociation. Formation of the plus-one intermediate was rate limiting for template switching. PV RdRp also utilized synthetic, preformed intermediates, including those with UMP 3’-overhangs. Reactions showed up to five consecutive template-switching events, consistent with a repair function for this form of recombination. PV RdRp may exclude UMP during forced-copy-choice RNA recombination to preclude creation of nonsense mutations during RNA fragment assembly. Several other picornaviral RdRps were evaluated, and all were capable of RNA fragment assembly to some extent. Lastly, we propose a structure-based hypothesis for the PV RdRp-plus-one intermediate complex based on an elongating PV RdRp structure.

## Introduction

A common property of most, if not all, positive-strand RNA viruses of bacteria, plants, and animals is the ability to recombine ^1, 2, 3^. Based primarily on mechanistic studies performed using poliovirus (PV), RNA recombination is thought to use a template-switching mechanism in which the RNA-dependent RNA polymerase (RdRp) initiates RNA synthesis on a *donor* template then switches to an *acceptor* template to complete RNA synthesis ^4^. This type of recombination is referred to as copy-choice recombination because the RdRp has a choice in the template that it copies (**Fig. 1A**). This template switch requires sequence complementarity between the 3’-end of nascent RNA and the acceptor template (**Fig. 1A**). When the acceptor template is a different RNA molecule than the donor template, an intermolecular recombinant is produced, carrying some genetic traits from both the donor and acceptor (**Fig. 1A**). In contrast, when the acceptor template is the same RNA molecule, an intramolecular recombinant is produced that harbors a deletion relative to the original donor template. This mechanism is at least one source of defective enteroviral genomes ^1, 5^. Both intermolecular and intramolecular recombination appear to be triggered by the same molecular mechanisms, including nucleotide misincorporation and incorporation of certain nucleotide analogs by the RdRp ^6–10^.

**Figure 1.**
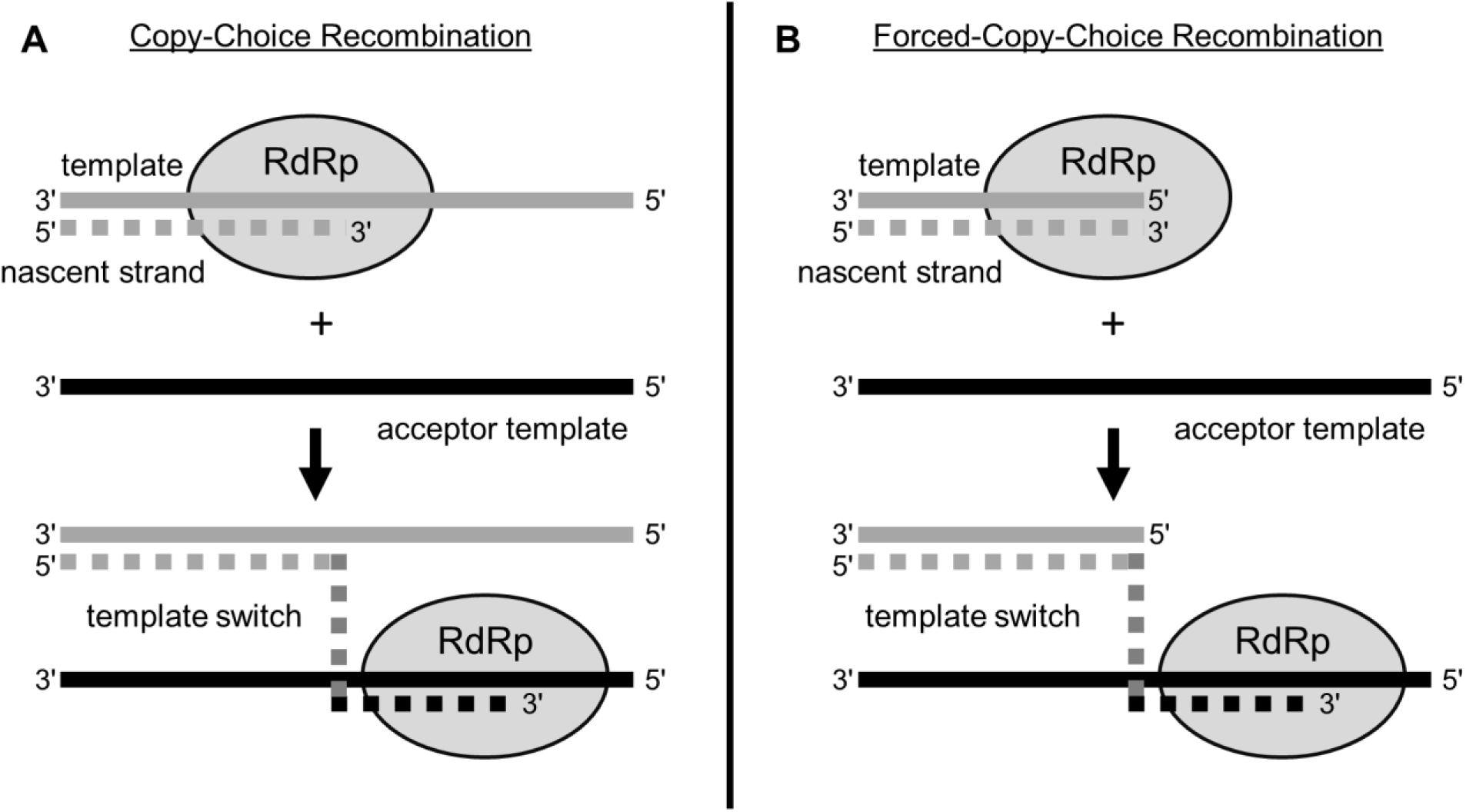
Viral RNA recombination can occur by either a copy-choice or a forced-copy choice mechanism. During viral RNA synthesis, the RdRp elongates the nascent RNA strand while copying a template strand and can subsequently switch templates by either a copy-choice or a forced-copy-choice mechanism. (**A**) Copy-choice recombination occurs with the RdRp switching templates at an internal position while continuing to extend the nascent RNA strand by copying the acceptor template. (**B**) Forced-copy-choice recombination occurs when the RdRp reaches the end of the initial template, for example at a premature end caused by a ribonuclease. The RdRp continues to extend the nascent RNA by switching templates and copying the acceptor template.

The selective pressure forcing viruses to maintain the capacity to recombine, particularly in cell culture, is unclear. PV mutants with defects to copy-choice RNA recombination can replicate as well or slightly better than wild-type virus in cell culture ^10–12, 13^. However, these viruses are highly attenuated in vivo ^11, 13, 14^. The function for copy-choice RNA recombination in the viral lifecycle and/or pathogenesis is also unclear ^3^. One possibility is that recombination purges accumulated, mutations caused by RNA synthesis by the error-prone RdRp ^11, 15^. One recent study from our group showed that copy-choice RNA recombination could be selectively impaired by mutations in RdRp-coding sequence, but this PV mutant exhibited impaired infection dynamics in cell culture relative to wild-type ^16^. The reason that other mutants failed to reveal such a phenotype may reflect the magnitude of the recombination defect, how recombination was measured, and/or impact on some as yet-to-be-defined property of the RdRp required for RNA recombination ^16^.

A second type of recombination is forced-copy-choice RNA recombination. This mechanism occurs when an elongating RdRp reaches the end of the donor template. This end is often not the authentic end of the genome; rather, it may have been created by nucleolytic cleavage of the donor template (**Fig. 1B**) ^17–19^. An obligatory step of reverse transcription used by retroviruses requires a forced-copy-choice mechanism ^18–21^. Reverse transcriptase (RT) initiates DNA synthesis using a tRNA primer annealed to the primer-binding site of the RNA genome at a site located a few hundred nucleotides or so from the 5’-end ^19, 20^. The product of this RT-catalyzed elongation reaction is a strong-stop DNA that uses complementarity between its 3’-end and that of the 3’-end of the genome in a *forced* template switch ^19–21^. This reaction has been mimicked *in vitro* using synthetic nucleic acids and purified RT from human immunodeficiency virus (HIV) ^6, 17, 22, 23^. A hallmark of this reaction *in vitro* was the addition of a non-templated nucleotide to the strong-stop DNA product that was present in the product of template switching ^7, 17, 23^. However, no function of the non-templated nucleotide was proposed, as the forced template switch was thought to require substantial complementarity between strong-stop DNA and acceptor template ^7, 17, 23^.

Since the description of the mechanism of HIV RT-catalyzed forced-copy-choice recombination, evidence has accumulated for the existence of a terminal transferase-like activity of RTs broadly that permits addition of non-templated nucleotides to the 3’-end of strong-stop DNA ^17, 21, 24^. Interestingly, the preferred nucleobase and number of nucleotides added vary ^17, 21, 24^. Most recently, studies of a group II intron-encoded RT have shown that basepairing between a single, non-templated nucleotide on the donor template and the 3’-end of an acceptor template is sufficient for template switching ^25^.

Enteroviruses initiate RNA synthesis using a VPg peptide ^26, 27^ or VPg-containing precursor ^28^ and a structured template located in protein 2C-coding sequence of the genome ^29^. The resulting di-uridylylated VPg must then switch to the 3’-end of the viral genome to produce full-length, negative-strand RNA. This process is akin to the first steps of retroviral reverse transcription described above. Efforts to mimic extensive, homology-driven, copy-choice RNA recombination using synthetic RNA templates and purified PV RdRp demonstrated template switching *in vitro* ^10, 30^. However, subsequent studies suggested forced-copy-choice RNA recombination as the likely mechanism observed ^10^.

Here, we report the development of a primed-template system that permits a more rigorous evaluation of the mechanism of forced-copy-choice RNA recombination catalyzed by PV RdRp. To our surprise, the mechanism used by PV RdRp and other picornaviral RdRps is strikingly similar to that described for the group II intron-encoded RT. We present a structure-based hypothesis for the mechanism of enteroviral RdRp-catalyzed, forced-copy-choice RNA recombination. Using this mechanism, enteroviral RdRps can assemble RNA fragments together randomly. We discuss the possibility of using this reaction as a last-ditch mechanism to rescue a full-length viral genome when faced with nucleolytic damage.

## Materials and Methods

### Materials

RNA oligonucleotides were from Horizon Discovery Ltd. (Dharmacon). T4 polynucleotide kinase was from ThermoFisher. [γ-^32^P]ATP (6,000 Ci/mmol) was from Perkin Elmer. Nucleoside 5’-triphosphates (ultrapure solutions) were from Cytiva. Heparin was from MilliporeSigma. All other reagents were of the highest grade available from MilliporeSigma, VWR, or Fisher Scientific.

### RNA purification

RNA substrates were suspended in 90 mM Tris, 90 mM Boric Acid, 2 mM EDTA, 90% formamide and resolved by denaturing polyacrylamide gel electrophoresis on an 18.5% acrylamide / 1.5 % bisacrylamide gel containing 7M Urea and 1X TBE. RNA bands were visualized by UV shadowing (260 nm) and excised from the gel. Excised gel samples were crushed and RNA was eluted by using an Elutrap Electroelution System (GE Healthcare). The RNA was concentrated by Sep-Pak C18 Classic Cartridge (Waters) and solvent was evaporated in a Speed Vac Concentrator (Savant). RNA samples were deprotected by adding 500 mM Acetic Acid and heating to 65 °C for 15 minutes, then adding 660 mM Tris base and heating to 65 °C again for 15 minutes. Final preparation of RNA samples involved desalting by running through G25 Sephadex equilibrated with 10 mM Tris pH 8.0, 1 mM EDTA (Sigma-Aldrich) column.

### 5’-32P-labeling of RNA substrates

RNA oligonucleotides were end-labeled by using [*γ*- ^32^P]ATP and T4 polynucleotide kinase. Reaction mixtures, with a typical volume of 50 *μ*L, contained 0.5 *μ*M [*γ*-^32^P]ATP, 10 *μ*M RNA oligonucleotide, 1X kinase buffer, and 0.4 unit/*μ*L T4 polynucleotide kinase. Reaction mixtures were incubated at 37 °C for 60 min and then held at 65 °C for 5 min to heat inactivate T4 PNK.

### Annealing of dsRNA substrates

dsRNA substrates were produced by annealing 10 *μ*M RNA oligonucleotides in T_10_E_1_ [10 mM Tris pH 8.0 1 mM EDTA] with 50 mM NaCl in a Progene Thermocycler (Techne). Annealing reaction mixtures were heated to 90 °C for 1 min and slowly cooled (5 °C/min) to 10 °C. Specific scaffolds are described in the figure legends.

### Expression and purification of PV RdRp

Expression and purification of WT and K359R PV RdRps were performed essentially as described previously ^31, 32^.

### Expression and purification of Picornaviral RdRps

Expression and purification of Picornaviral RdRps (CVB3, EV-D68, RV-C15, RV-A16, and FMDV) were performed essentially as described previously ^33^. The following accession numbers are provided as reference sequences for the indicated Picornaviral RdRps: CVB3: JX312064.1; EV-D68: KT347249.1; RV-C15: GU219984; RV-A16: L24917.1; and FMDV: AJ133357.1.

### *In vitro* RdRp-primed-template complex assembly assays

Elongation complexes were assembled by incubating 5 µM WT or mutant poliovirus polymerase with 0.5 µM RNA primed-template duplex and 500 µM ATP for various amounts of time before being quenched by addition of 50 mM EDTA. Reactions included heparin (0 to 60 µM) during assembly or after assembly of enzyme onto RNA duplex. All and subsequent reactions described were performed at 30°C in 50 mM HEPES pH 7.5, 10 mM 2-mercaptoethanol, 5 mM MgCl_2_ and 60 µM ZnCl_2_.

### Complex stability assay

Elongation complexes were assembled by incubating 5 µM WT or mutant poliovirus polymerase with 0.5 µM RNA primed-template duplex and 500 µM ATP. At various time points, 500 µM UTP was added and the reaction was quenched 30 seconds later by addition of 50 mM EDTA.

### *In vitro* template-switching assay

Elongation complexes were assembled by incubating 5 µM WT or mutant poliovirus polymerase with 0.5 µM RNA primed-template duplex and 500 µM ATP for 30 minutes (Mix 1). Template-switching reactions were initiated by adding 60 µM of RNA acceptor template and 500 µM CTP, GTP, UTP and 25 µM heparin (Mix 2) and quenched at the listed times by addition of 50 mM EDTA.

### Denaturing PAGE analysis of reaction products

An equal volume of loading buffer (85% formamide, 0.025% bromophenol blue and 0.025% xylene cyanol) was added to quenched reaction mixtures and heated to 90 °C for 5 min prior to loading 5 µL on a denaturing 23% polyacrylamide gel containing 1X TBE (89 mM Tris base, 89 mM boric acid, and 2 mM EDTA) and 7 M urea. Electrophoresis was performed in 1X TBE at 90 W. Gels were visualized by using a PhosphorImager (GE) and quantified by using ImageQuant TL software (GE).

### Sanger sequencing and small RNA-Seq of RNA products

In vitro template switching assays were performed with 5’-phosphate-terminated primers and 3’-deoxy-terminated acceptor templates. Reactions were quenched with EDTA after 60 min, reaction products were resolved by denaturing PAGE and both strong-stop and transfer products were gel purified. RNA products were either cloned and sequenced following the small RNA cloning kit miRCat-33^TM^ protocol (Integrated DNA Technologies) or total purified RNA was submitted for small RNA-Seq next generation sequencing (Illumina, 2×150 bp, ∼350M PE reads, single index) performed by Genewiz. Sanger sequencing results (> 100 sequences) were aligned to the following reference primer and transfer product sequence (5’-UUUUCCGGGC-3’; 5’- UUUUCCGGGCAUGCCCAUGCAUGCUUGC-3’). Results from NGS used the following pipeline for analysis: data was evaluated with FastQC, trimmed with bbduk.sh (BBmap suite) to remove adapter sequences (both Illumina universal and small RNA adapter), sequences were clustered using clumpify.sh (BBMap suite), each sequence was treated differently which did not allow any substitutions, and the top 500 sequences identified (sequence, count, and length).

### Data analysis

All gels shown are representative, single experiments that have been performed at least three to four individual times to define the concentration or time range shown with similar results. In all cases, values for parameters measured during individual trials were within the limits of the error reported for the final experiments. Data were fit by either linear or nonlinear regression using the program GraphPad Prism v7.03 (GraphPad Software Inc.).

## Results

### New primed-templates to study template switching by the enterovirus RdRp

When our laboratory began studying PV RdRp, we pursued the use of conventional primed-templates: a stable duplex with a 10-nt or longer 5’ overhang to serve as template ^34^. To our surprise, the RdRp partitioned in an orientation such that it was bound to the duplex portion of the substrate instead of at the primed-template junction ^35^. When bound to the duplex, the enzyme was able to add one or more non-templated nucleotides depending on the divalent cation used in the reaction ^35^. These observations led to development of a symmetrical, primed-template (sym/sub) that had a duplex formed by a self-complementary sequence with 5’-overhangs on both strands ^34^. This substrate proved to be the first that could be used for pre-steady-state kinetic analysis of PV RdRp-catalyzed nucleotide incorporation ^36^. Unfortunately, using this substrate to study template switching is complicated by the fact that one cannot distinguish primer from template because of the self-complementary nature of the RNA used.

To address this problem, we have created primed-templates that are no longer self-complementary (**Fig. 2A**). To the 5’-end of the primer (P_0_) and 3’-end of template (T), we added four uridine nucleotides to prevent the enzyme from binding in an unproductive conformation, although the addition to the primer turned out not to be essential (**Fig. S1**). Assembly of PV RdRp on this primed-template to form a stable elongation complex was slow (t_1/2_ = 7 min) (**Figs. 2B-2D**), but the resulting elongation complex was very stable (t_1/2_ = 540 min) (**Figs. 2E-2G**).

**Figure 2.**
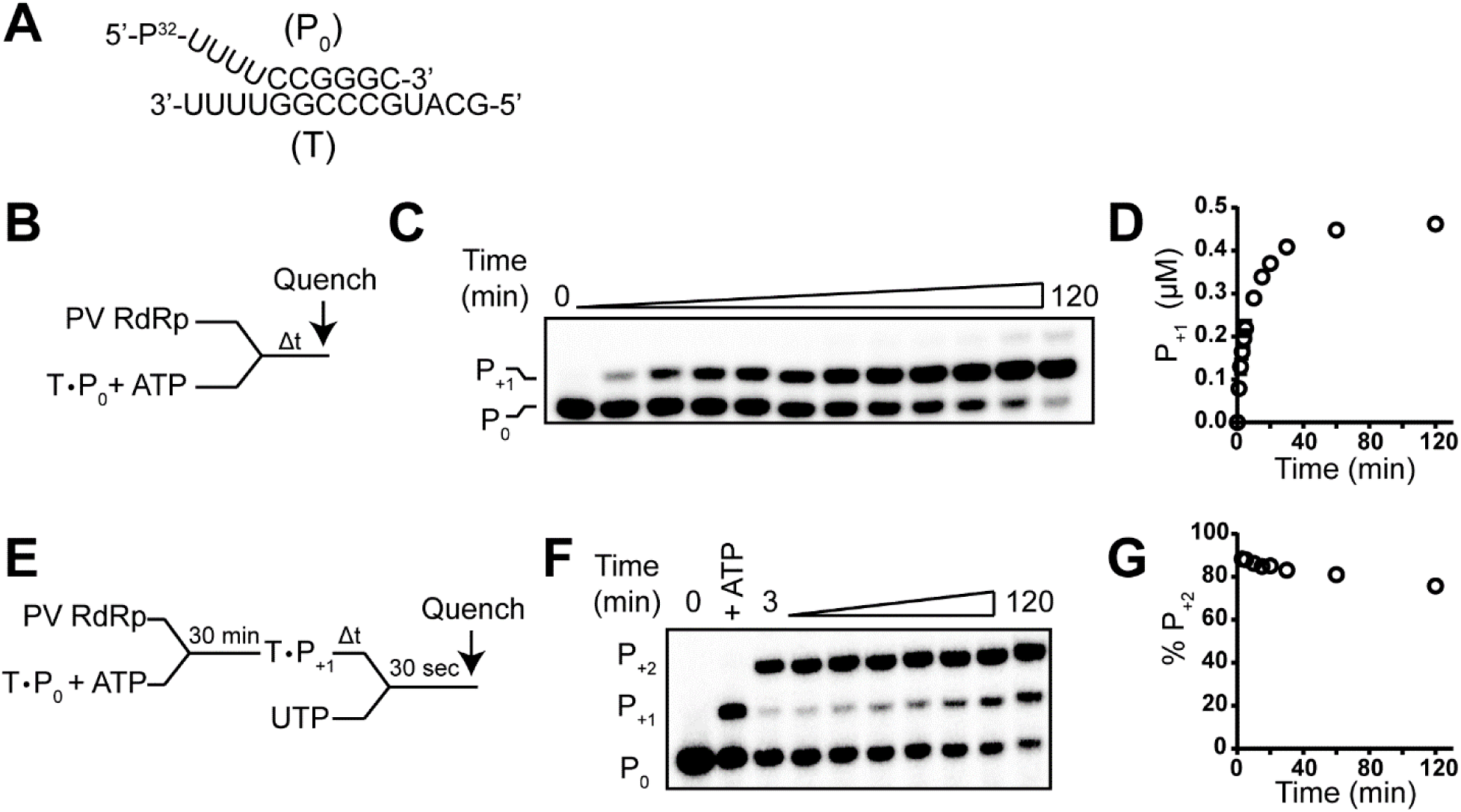
Establishment of a primed-template to study template switching by PV RdRp. (**A**) Primed-template used in this study. The primer (P_0_) is a 10-nt RNA and template (T) is a 14-nt RNA; sequences are shown. The annealed primed-template forms a 6-bp duplex with a 4-nt 5’-template overhang. The RNA primer was labeled on the 5’-end with ^32^P. (**B**) Schematic of assay to measure assembly of the elongation complex. Primer extension was initiated by adding PV RdRp with primed-template in the presence of ATP for various amounts of time and then quenched. (**C**) Analysis of reaction products by denaturing PAGE. The positions of the unextended primer P_0_ and extended primer P_+1_ are indicated. (**D**) Quantitative analysis of the kinetics of assembly by monitoring product RNA (P_+1_) formation. (**E**) Schematic of assay to measure stability of the elongation complex. Elongation complexes were assembled by adding PV RdRp with primed-template in the presence of ATP for 30 min. After formation of P_+1_, UTP was added after various amounts of time and reactions quenched after 30 s. (**F**) Analysis of reaction products by denaturing PAGE. The positions of the unextended primer P_0_ and extended primers P_+1_ and P_+2_ are indicated. (**D**) Quantitative analysis of elongation complex stability over time by monitoring the percentage of product RNA (P_+2_) formed relative to total product RNA (P_+1_ + P_+2_).

To this primed-template we added an acceptor template (T_A_) that was used previously with sym/sub ^10^. The acceptor template was designed to study copy-choice RNA recombination, with two, sequential sites that would be complementary to the product produced by elongation of the primer (**Fig. 3A**). Later, it became clear that, while this system supported template switching, it was not mimicking copy-choice RNA recombination ^10^. Rather, it was mimicking forced-copy-choice RNA recombination ^10^.

**Figure 3.**
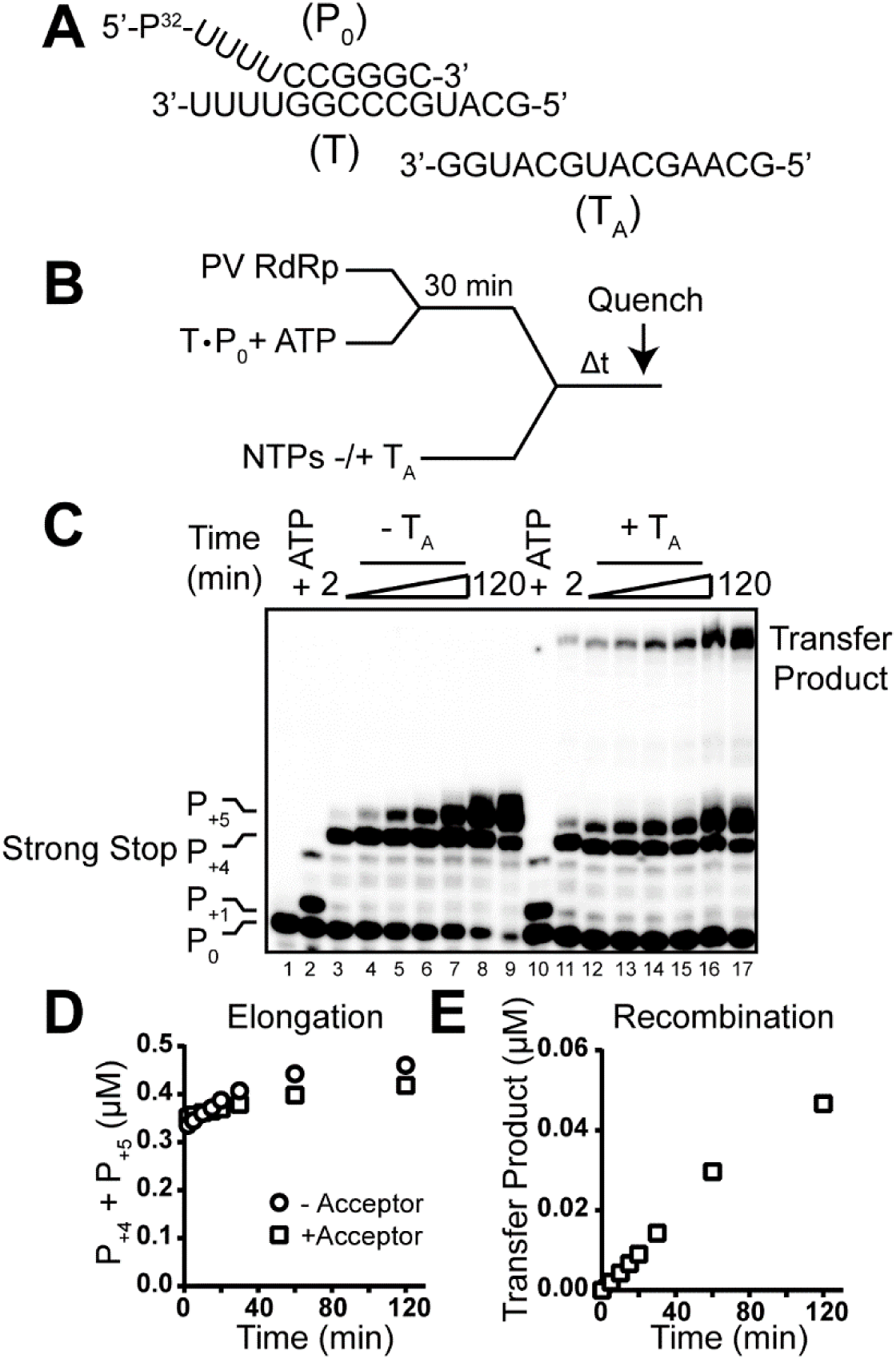
PV RdRp template-switching assay. (**A**) Primed-template and acceptor template (T_A_) RNAs used to assess template switching by PV RdRp. (**B**) Schematic of assay used to measure template switching. Elongation complexes were assembled by adding PV RdRp with primed-template in the presence of ATP for 30 min at which point acceptor template and all four NTPs were added. Reactions were allowed to proceed for various amounts of time and then quenched. (**C**) Analysis of reaction products from reactions performed in the absence or presence of acceptor template by denaturing PAGE. The positions of the unextended primer (P_0_), extended primer (P_+1_), strong-stop product (P_+4_), strong-stop product with the addition of a non-templated nucleotide (P_+5_), and transfer product are indicated. The transfer product is only observed in reactions performed in the presence of acceptor template (T_A_). (**D,E**) Quantitative analysis of the formation of strong-stop (P_+4_ + P_+5_) products (Elongation) or transfer product (Recombination) as function of time. Mean of three replicates are shown.

We mixed RdRp, primed-template, and ATP (first templated nucleotide) for a time sufficient to produce a one-nucleotide-extended product (P_+1_) (**Fig. 3B**) whose fate could be monitored (lane 2 of **Fig. 3C**). In the presence of all four nucleotides but the absence of acceptor template, two major products formed (lanes 2-9 of **Fig. 3C**). The first product was consistent with the RdRp elongating to the end of template; we refer to this product as strong-stop or P_+4_ (**Fig. 3C**). Over time, the strong-stop product was extended by one nucleotide in a non-templated fashion; we refer to this product as the product of non-templated addition, the “plus-one” product, or P_+5_ (**Fig. 3C**). In the presence of the acceptor template, we observed a product of template switching, referred to as the transfer product (**Fig. 3C**). Elongation was equally efficient in the absence and presence of acceptor template (**Fig. 3D**). Template switching occurred at a rate of 4 ×10^−3^ µM/min using 5 µM of PV RdRP (**Fig. 3E**).

To determine if template switching occurred processively—that is, without dissociation of the enzyme used to produce the strong-strop and/or plus-one products, we required a trap for free or dissociating enzyme. Using sym/sub, heparin worked well ^10, 30^. With the new substrate, heparin showed complete inhibition at 20 µM (**Figs. 4A-4B**). When we formed the elongation complex and then added heparin along with the remaining three nucleotides, heparin exhibited no effect on the production of strong-stop or plus-one products (**Figs. 4C-4D**). We repeated this experiment in the presence of the acceptor template (**Fig. 5**). Again, neither formation nor utilization of the elongation complex was impaired in the presence of heparin (**Figs. 5B-5C**). While production of the transfer product was not completely resistant to heparin, at least 20% of the complexes were (**Fig. 5D**).

**Figure 4.**
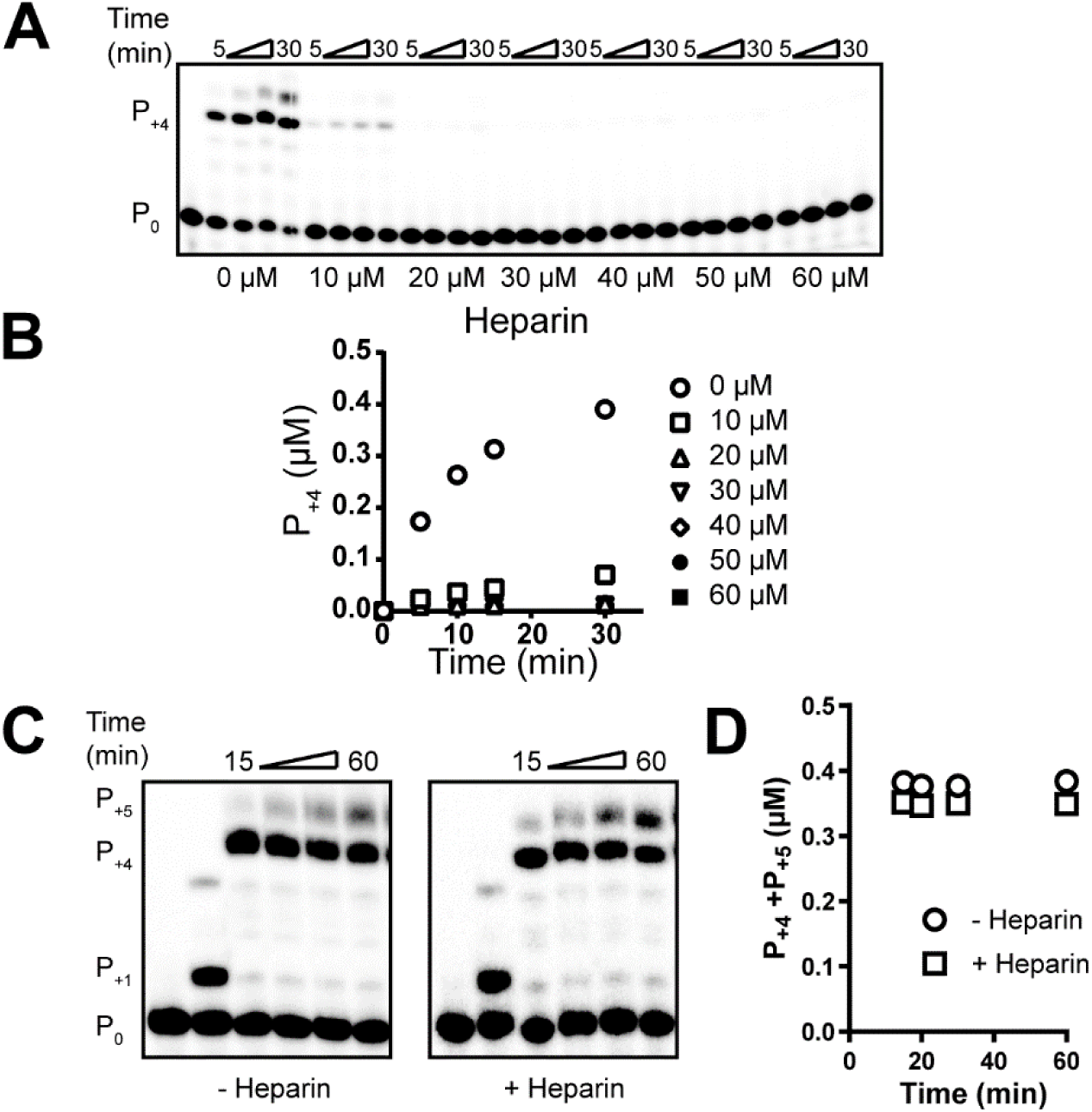
Heparin inhibits complex assembly but not elongation. (**A**) Complex assembly in the presence of increasing concentrations of heparin. Primer extension was initiated by adding PV RdRp with primed-template in the presence of all four NTPs and the indicated amounts heparin for 5 to 30 min and then quenched. Shown is the analysis of reaction products resolved by denaturing PAGE. (**B**) Quantitative analysis of the kinetics of assembly by monitoring product RNA (P_+4_) formation in the presence of increasing concentrations of heparin. (**C**) Elongation in the presence of heparin. Elongation complexes were assembled by adding PV RdRp with primed-template in the presence of ATP for 30 min. After formation of P_+1_, all four NTPs with or without heparin were added and reactions quenched. Shown is the analysis of reaction products resolved by denaturing PAGE. (**D**) Quantitative analysis of the kinetics of elongation by monitoring product RNA (P_+4_) formation in the presence of heparin.

**Figure 5.**
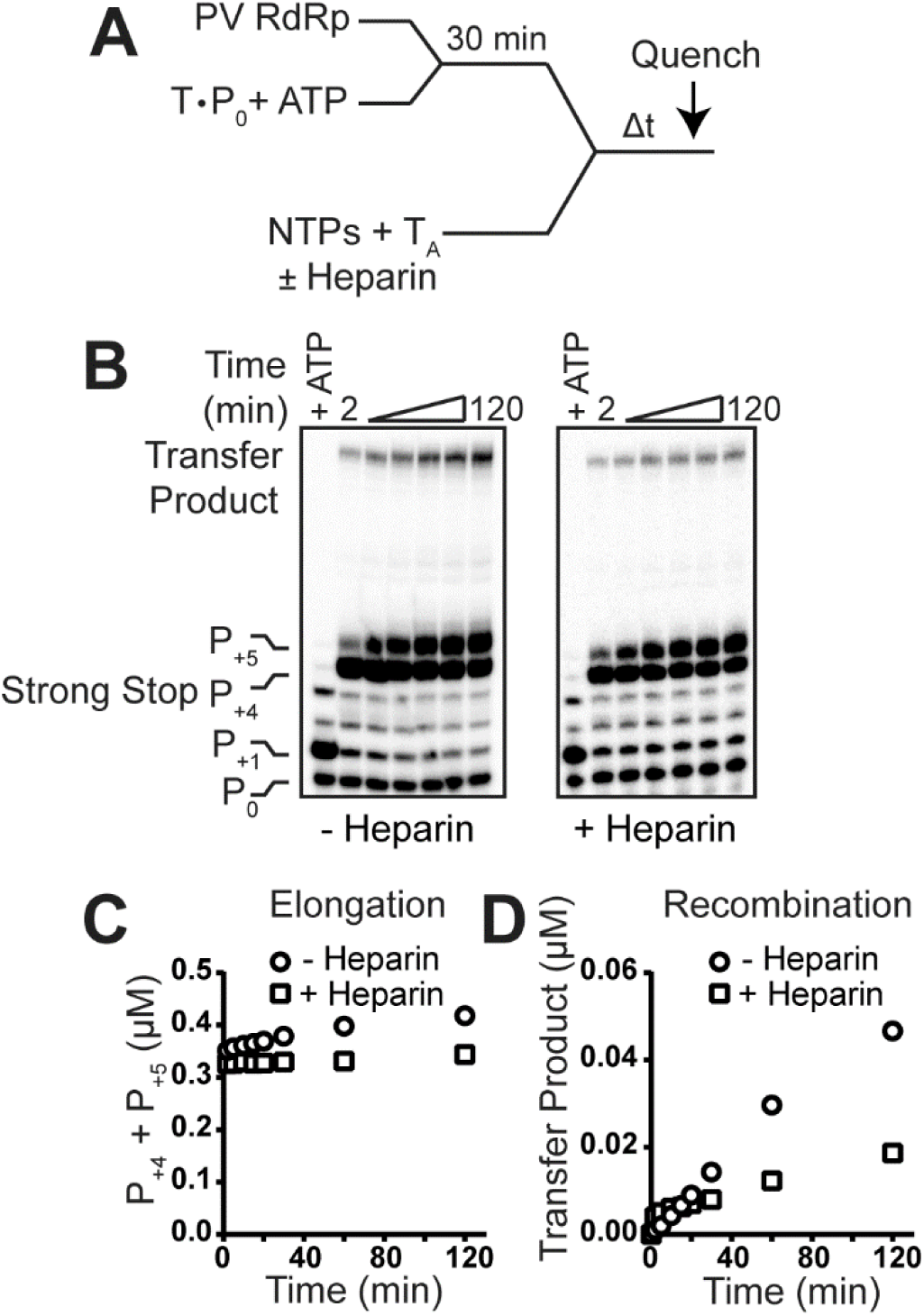
PV RdRp-catalyzed template switching can occur by a heparin-resistant mechanism. (**A**) Schematic of template-switching assay performed in the absence or presence of heparin. Elongation complexes were assembled by adding PV RdRp with primed-template in the presence of ATP for 30 min at which point acceptor template and all four NTPs were added in the presence or absence of heparin. Reactions were allowed to proceed for the indicated amount of time and then quenched. (**B**) Analysis of reaction products by denaturing PAGE from template-switching reactions performed in the absence or presence of heparin. The positions of the unextended primer (P_0_), extended primer (P_+1_), strong-stop product (P_+4_), strong-stop, “plus-one” product (P_+5_), and transfer product are indicated. The transfer product is observed in reactions performed in the presence of heparin. (**C,D**) Quantitative analysis of the formation of strong-stop products or transfer-product as function of time in the absence or presence of heparin. The total amount of transfer product in the absence and presence of heparin at 120 min was 0.047 µM and 0.019 µM, respectively.

### The plus-one product as an intermediate required for forced-copy-choice RNA recombination

In order to gain insight into how the transfer product was formed, we cloned and sequenced the strong-stop, plus-one, and transfer products from reactions performed under a variety of conditions, including in the absence and presence of heparin, as described under Materials and Methods (**Table 1**). As expected from the gel-based assays, the sequence(s) observed for strong-stop and plus-one products were as expected, and the presence of acceptor template did not alter the sequence(s) observed (compare section A to section B in **Table 1**). In the presence of all four ribonucleoside triphosphates, each could be used for non-templated addition. Addition of CMP and GMP were most efficient. Addition of AMP was easily detected. We only observed one example of UMP addition (sections A and B of **Table 1**). The sequence of the transfer product could only be explained as follows: (1) strong-stop RNA is produced; (2) non-templated additions occur, producing plus-one products; (3) the plus-one product capable of basepairing to the 3’-end of acceptor template is selected; and (4) continued synthesis leads to production of a 28-nt RNA transfer product (sections C and D of Table 1). Therefore, we suggest that the plus-one product is an obligatory intermediate for forced-copy choice RNA recombination (**Fig. 6A**). Use of the plus-CMP product was preferred in the absence and presence of heparin (compare section C to section D in **Table 1**). We observed a few other transfer-product sequences that may have arisen from misincorporation or utilization of a truncated template during strong-stop RNA synthesis (see lines 2 and 3 of sections C and D of **Table 1**).

**Figure 6.**
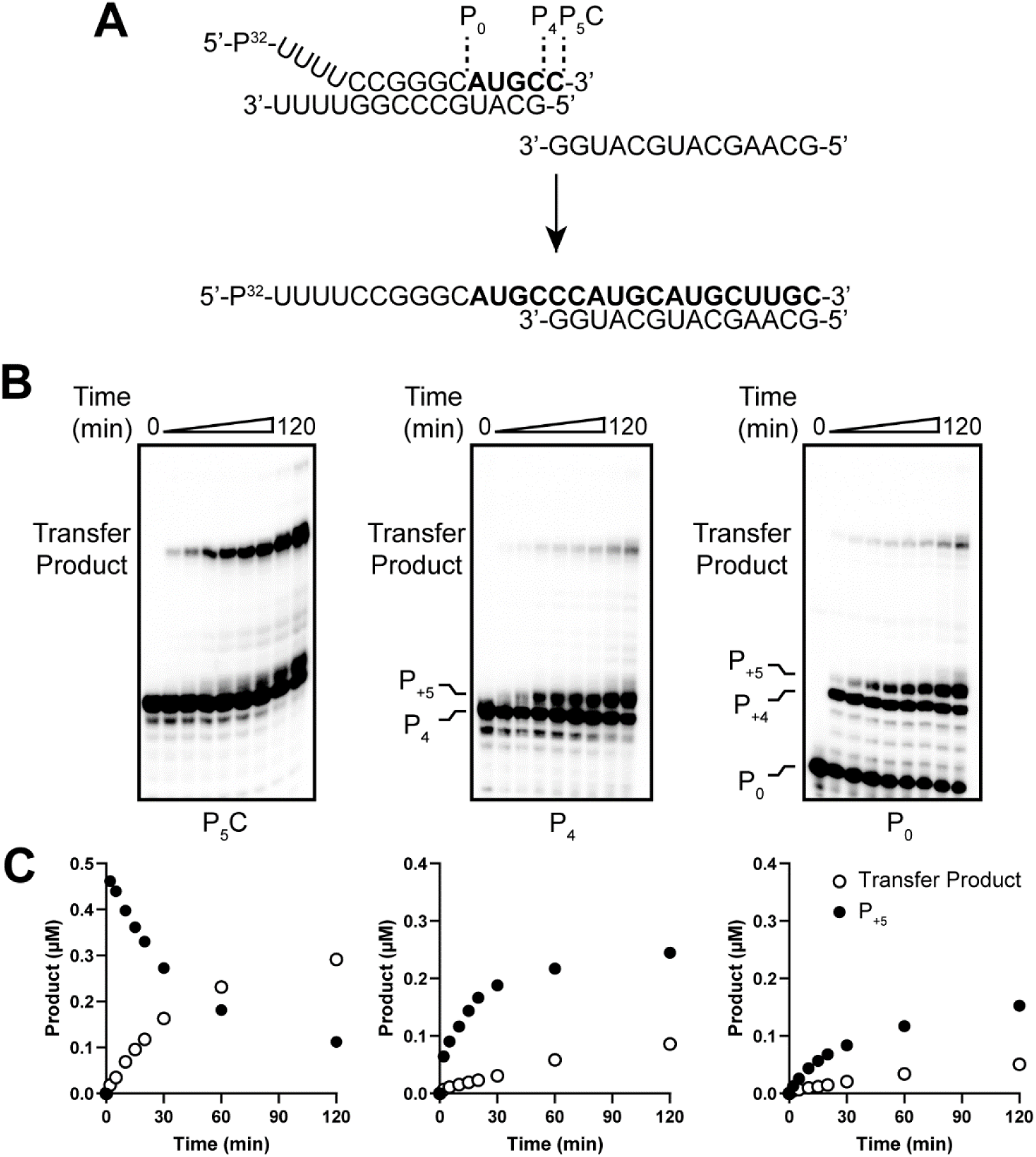
The plus-one product (P_+5_) as an intermediate in forced-copy-choice recombination. (**A**) Primers of varying length (P_0_, P_4_, and P_5_C) used to assess template switching by PV RdRp. P_4_ and P_5_C are consistent with strong-stop products produced during elongation: blunt-ended (P_4_) or 3’-overhang formed by non-templated addition (“plus-one” product, P_5_C). (**B**) Analysis of reaction products by denaturing PAGE from template-switching reactions performed with P_0_, P_4_, and P_5_Cprimed-template duplexes. The positions of the unextended primer, strong-stop product (P_+4_), strong-stop, “plus-one” product (P_+5_), and transfer product are indicated. (**C**) Quantitative analysis of the formation of transfer product as function of time from template-switching reactions performed with P_0_, P_4_, and P_5_C primed-template duplexes. Reactions using P_5_C primed-template duplexes resulted in transfer products six-fold greater than reactions with P_0_ or P_4_. Mean of three replicates are shown. Error bars represent standard deviation.

**Table 1.**
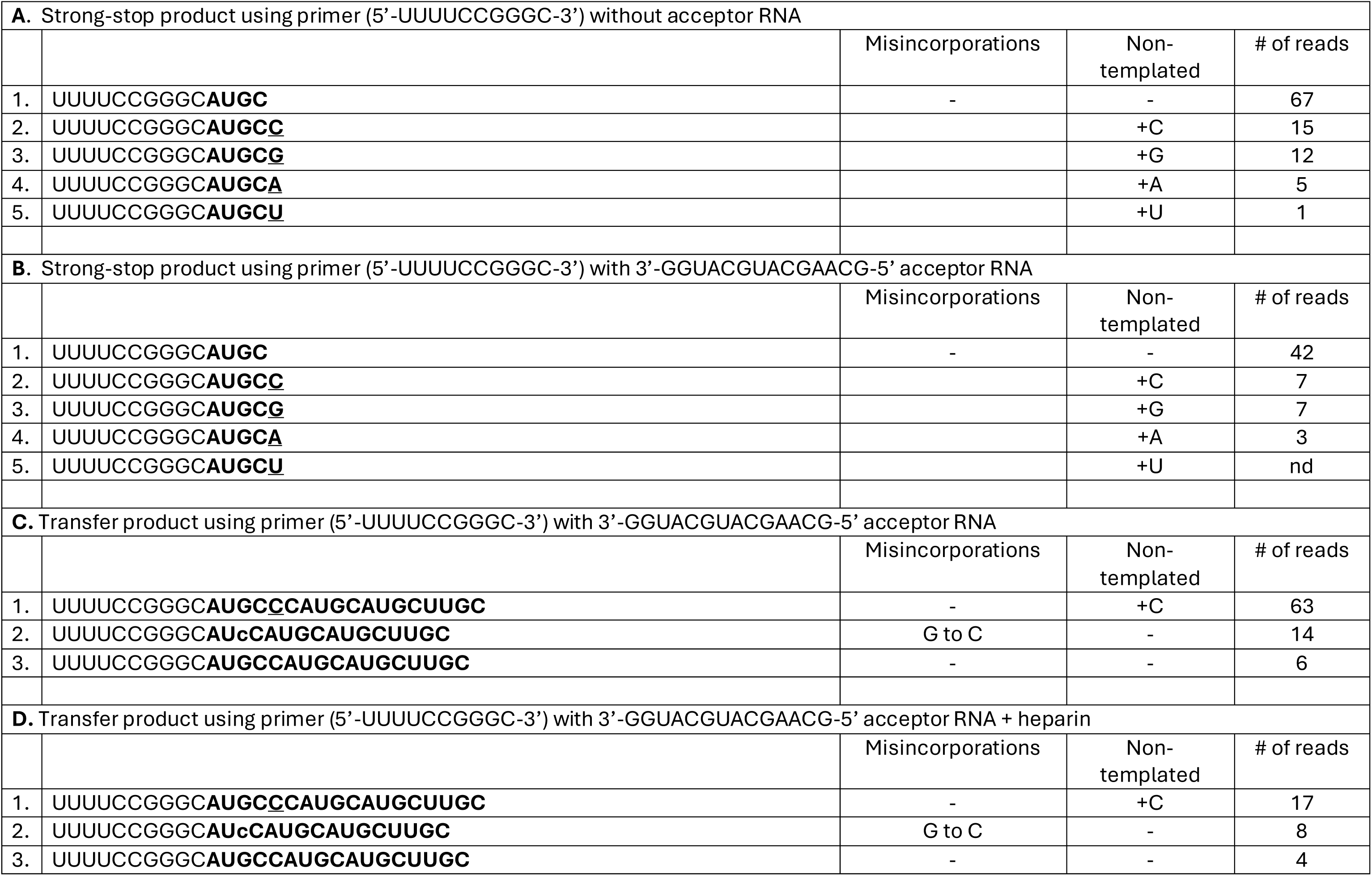
Sequence analysis of strong-stop and transfer products. The last column indicates the total number of each sequence obtained. Bold nucleotides indicate extension of primer. Underlined nucleotides indicate non-templated nucleotide addition and lower case indicates misincorporation (identified explicitly in the indicated columns). nd: not detected

Isolation of RNA products from gels, cloning, and then sequencing clearly can introduce bias into the results. Therefore, we pursued direct analysis of RNA substrates, intermediates, and products from reactions performed in vitro using small RNA-Seq as described under Materials and Methods. The first observation was that even analysis of just the primer alone revealed a lot of background. We observed 500 different sequences in the preparation of purified primer defined from 10^7^ reads; only five million or 50% of those represented the sequence ordered (see **Table S1**). Given this complexity, we only present data for sequences whose origin can be explained. The incorporation of the first, correct nucleotide (AMP) was quite faithful, representing 82% of primers extended by one nucleotide (line 3 of **Table 2**). GMP, CMP, and UMP misincorporation occurred at a frequency of 11%, 6%, and 1%, respectively (lines 4-6 of **Table 2**). We observed strong-stop RNA product of the correct sequence, which represented 84% of recognizable 14-nt sequences (line 7 of **Table 2**). Others had a misincorporation (lines 8-9 of **Table 2**). This analysis revealed a strong, unexpected bias for the addition of non-templated nucleotides. Non-templated addition of CMP was preferred (79% of events), and non-templated addition of UMP was below the limit of detection (less than 1% of events) (line 10 in **Table 2**). Non-templated addition of AMP (20%) and GMP (1%) was observed (lines 11 and 12 of **Table 2**). Finally, in addition to the expected 28-nt transfer product (33% of total), we also observed a 20-nt transfer product that accumulated (67% of total).

**Table 2.**
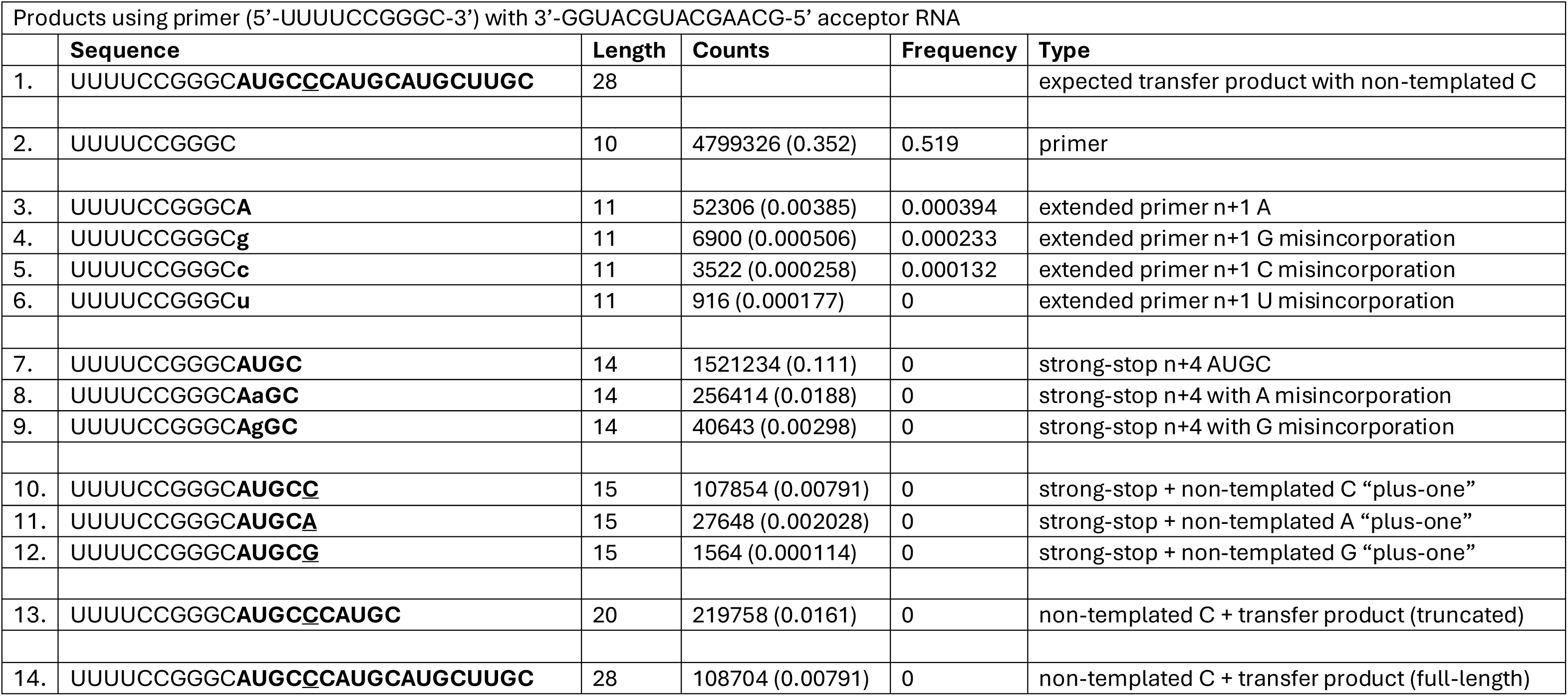
Sequence analysis of RNA products from small RNA-Seq. Representative sequences that correspond to primer, extended primer, strong-stop, “plus-one”, and transfer products, both truncated and full-length, are listed. The list is sorted from shortest to longest sequence. The sequence, length, counts, frequency, and type are indicated. Bold nucleotides indicate extension of primer. Underlined nucleotides indicate non-templated nucleotide addition and lower case indicates misincorporation. The top 500 sequences are reported in supplemental data tables: NGS_3A and NGS_3A_Control. All T’s in the sequence in supplemental tables have been converted to U’s as reported in the Table below.

**Table 3.**
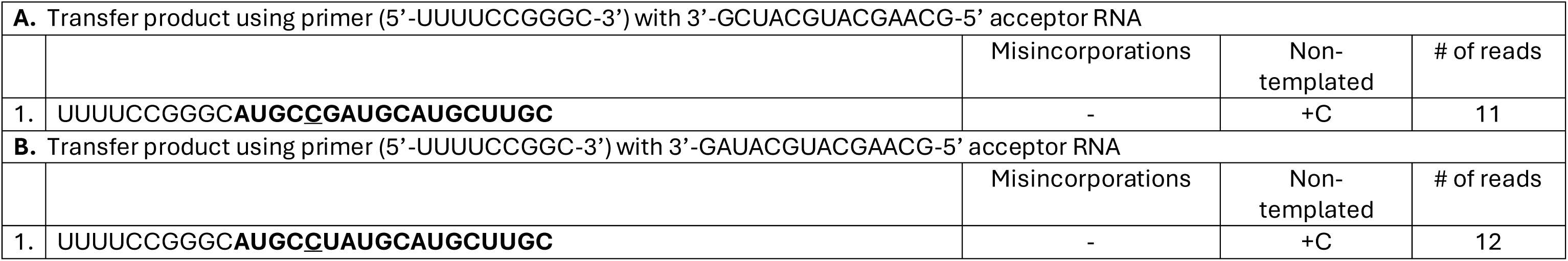
Sequence analysis of transfer products using different acceptors. The last column indicates the total number of each sequence obtained. Bold nucleotides indicate extension of primer. Underlined nucleotides indicate non-templated nucleotide addition and lower case indicates misincorporation (identified explicitly in the indicated columns). nd: not detected

The structure of the intermediate formed and capable of using a 3’-GMP-terminated acceptor template is shown in **Fig. 6A**. We asked if this intermediate could be used directly by PV RdRp in presence of acceptor to form a transfer product. PV RdRp used the plus-one intermediate (P_5_C in **Fig. 6B**) without delay. A P_+6_ product was also observed; however, it is unclear whether this product derived from a templated or non-templated event (**Fig. 6B**). In contrast, the formation of transfer product by utilization of a P_+4_ blunt-ended product that was preformed (P_4_ in **Fig. 6C**) or formed by extension of the P_0_ primer was limited by the rate of formation of the P_+5_ product. These data are consistent with formation of the plus-one intermediate as the rate-limiting step for PV RdRp-catalyzed forced-copy-choice RNA recombination.

Sequencing of the substrates and products produced during the reaction employing the P_5_C plus-one intermediate revealed accumulation of products representing each cycle of nucleotide addition, suggesting that translocation on the acceptor template differs from a template present in a conventional primed template (**Table 2**). The sequencing also revealed the error-prone nature of the PV RdRp during the template-switching process. We observed numerous misincorporation and deletion events (**Table 2**). Interestingly, we did not observe any insertions.

### The connection between RdRp nucleotide-incorporation fidelity and forced-copy-choice RNA recombination

The requirement for a plus-one intermediate for forced-copy-choice RNA recombination to occur provided a potential explanation for the inability of high-fidelity RdRp variants to support this type of RNA recombination ^10^. To test this possibility, we used a well-characterized, high-fidelity PV RdRp derivative, K359R ^31^. This derivative assembled and elongated primed templates as well as WT (**Figs. 7A** and **7B**) ^31^. However, unlike WT RdRp, K359R RdRp produced very little P_+5_ product after reaching the end of template (panel P_0_ in **Fig. 7C**). If production of the plus-one intermediate is the only defect for the high-fidelity RdRp, then the K359R RdRp should use the P_5_C intermediate. K359R RdRp used this intermediate to produce transfer product (panel P_5_C in **Fig. 7C**). Importantly, the rate of transfer product accumulation was equivalent to WT RdRp (**Fig. 7D**), consistent with the inability of K359R RdRp to produce the plus-one intermediate being the sole defect associated with the failure of this derivative to support forced-copy-choice RNA recombination.

**Figure 7.**
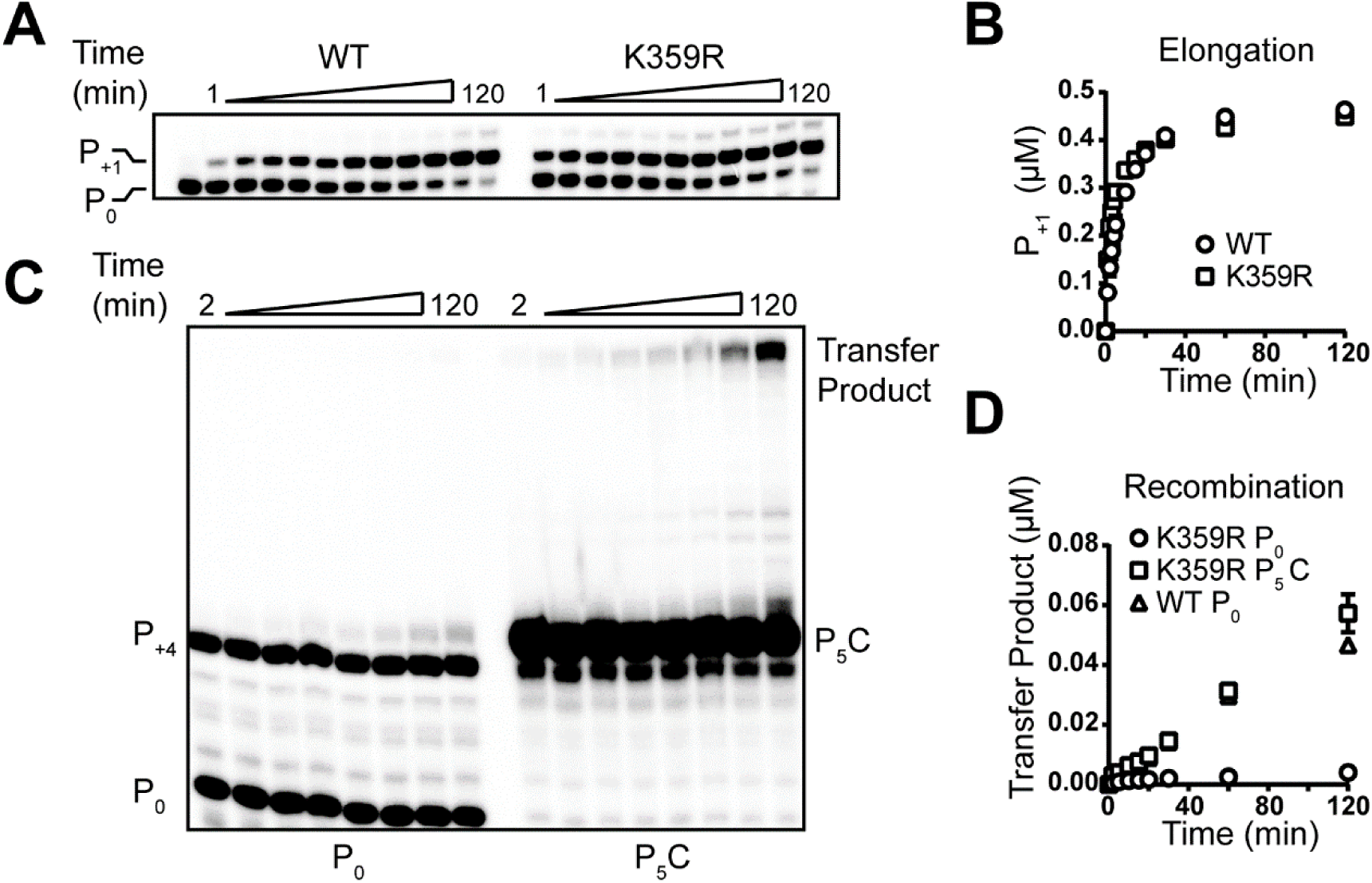
A high-fidelity polymerase derivative fails to catalyze forced-copy-choice recombination because of its inability to produce the plus-one intermediate. (**A**) Comparison of complex assembly between WT and K359R PV RdRp by monitoring primer-extension activity, formation of P_+1_ RNA product. Shown are the reaction products resolved by denaturing PAGE. (**B**) Quantitative analysis of the kinetics of assembly by monitoring product RNA (P_+1_) formation using either WT or K359R PV RdRp. There are no differences in the kinetics of assembly. (**C**) Analysis of reaction products by denaturing PAGE from template-switching reactions performed with P_0_ or P_5_C primed-template duplexes using K359R PV RdRp. The positions of the unextended primers (P_0_ or P_5_C), strong-stop product (P_+4_), and transfer product are indicated. Transfer product is only observed in reactions that used P_5_C-primed-template duplexes. (**D**) Quantitative analysis of the formation of transfer product as function of time from template-switching reactions performed using P_0_ and P_5_C primers with K359R PV RdRp and P_0_ primers with WT PV RdRp.

### Determinants of the intermediate required for efficient forced-copy-choice RNA recombination

We performed a series of experiments with modified intermediates and acceptor templates to identify determinants of each promoting template switching by PV RdRp.

#### Only a single, non-templated addition supports template switching

We synthesized an intermediate containing a two-nucleotide 3’-overhang (P_6_CC in **Fig. 8A**). Addition of this extra nucleotide completely inhibited formation of transfer product (**Fig. 8B**). Please note that the exposure time used was increased to maximize sensitivity. The band that appears above the primary, labeled primer in these and subsequent experiments represent no more than 10% of the total and likely derives from incomplete deprotection of the synthetic RNA. This band was observed in all images below derived from long exposures.

**Figure 8.**
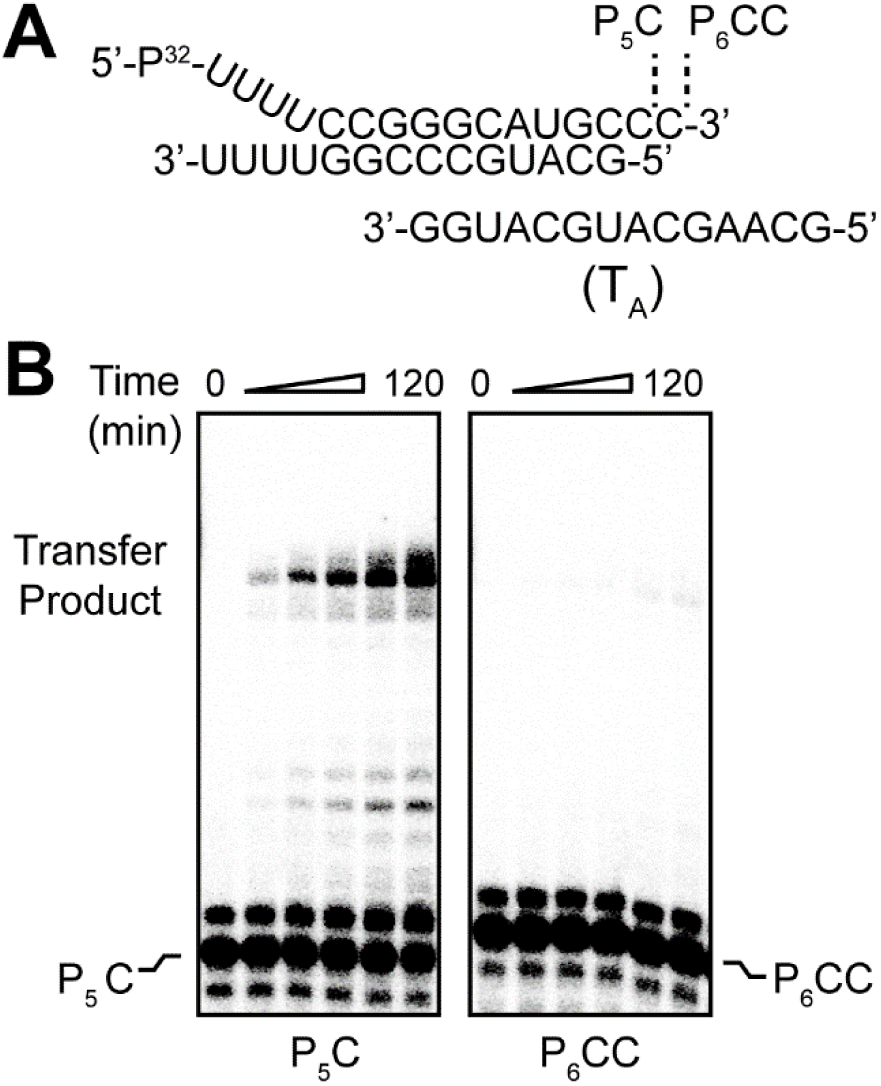
Only a single, non-templated nucleotide supports forced-copy-choice RNA recombination. (**A**) Duplexes containing 3’-overhangs of one (P_5_C) or two (P_6_CC) nucleotides were used to assess template switching by PV RdRp. (**B**) Analysis of reaction products by denaturing PAGE from template-switching reactions performed with P_5_C and P_6_CC primed-template duplexes. The positions of the unextended primers (P_5_C and P_6_CC) and transfer product are indicated.

#### Basepairing between the intermediate and acceptor template is required for template switching

To determine the importance of basepairing for acceptor template utilization, we created a terminal C:A mispair (**Fig. 9A**). This acceptor template failed to produce transfer product (**Fig. 9B**). We followed up this experiment with a more systematic analysis of the sufficiency of a single basepair for template switching. We created intermediates with each 3’-overhangs containing a different nucleotide (**Fig. 10A**). We then combined each intermediate individually with acceptor templates containing a different, terminal nucleotide (**Fig. 10A**). Only intermediates and acceptor templates with complementary nucleotides yielded transfer product (**Fig. 10B**). In all cases, transfer product accumulated over the two-hour period monitored; however, the efficiency of each reaction varied (**Fig. 10C**). While a C:G (intermediate:acceptor) basepair was most efficient (compare panel P_5_C in **Fig. 10C** to other panels), the G:C basepair was least efficient (compare panel P_5_G in **Fig. 10C** to other panels). The molecular basis for the observed differences in efficiency remains unclear and requires further investigation. By changing the context in which the terminal nucleotides of the intermediate and acceptor template were presented, we observed context-dependent changes in the efficiency of production of transfer product (**Fig. 11**). However, these changes were subtle (**Fig. 11**). Finally, we asked if the ribose hydroxyls of the terminal nucleotide of the acceptor template contributed to the efficiency of acceptor template utilization (**Fig. 12A**). We found that neither the 3’-OH nor the 2’-OH were required (**Fig. 12B**). Both the 3’-dG-terminated and the 2’-dG-terminated acceptor templates supported production of transfer product with the same efficiency as the 3’-G-terminated acceptor template when the intermediate was produced during the reaction (panels P_0_ in **Fig. 12B**) or the intermediate was produced synthetically (panels P_5_C in **Fig. 12B**).

**Figure 9.**
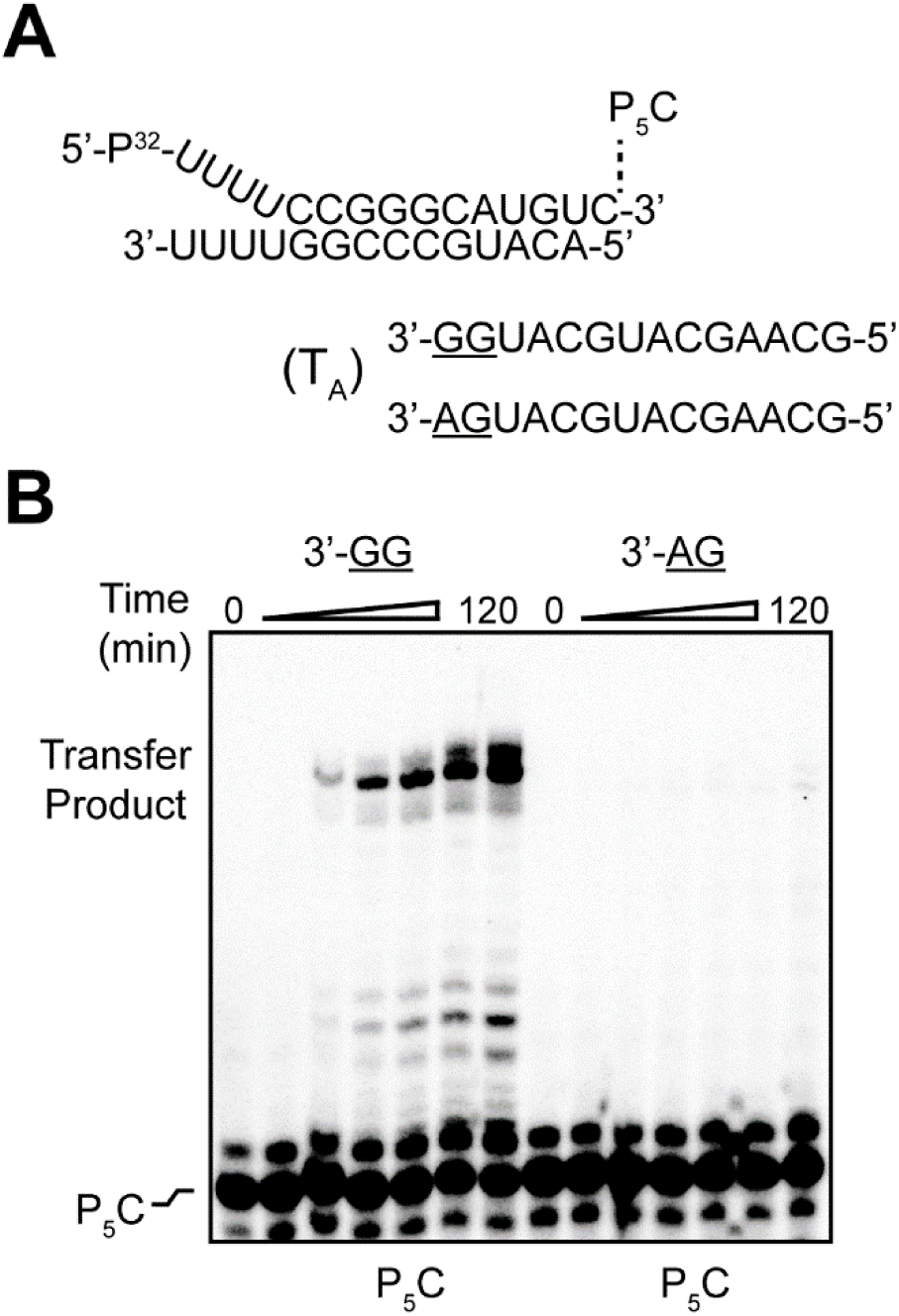
Basepairing to the 3’-end of the acceptor RNA is required for template switching. (**A**) P_5_C intermediate and acceptor template RNAs with unique 3’-terminal ends (3’-GG or 3’-AG end) used to assess template switching by PV RdRp. (**B**) Analysis of reaction products by denaturing PAGE from template-switching reactions performed with P_5_C primed-template duplex and 3’-GG or 3’-AG acceptor template RNA. The positions of the unextended primer and transfer product are indicated.

**Figure 10.**
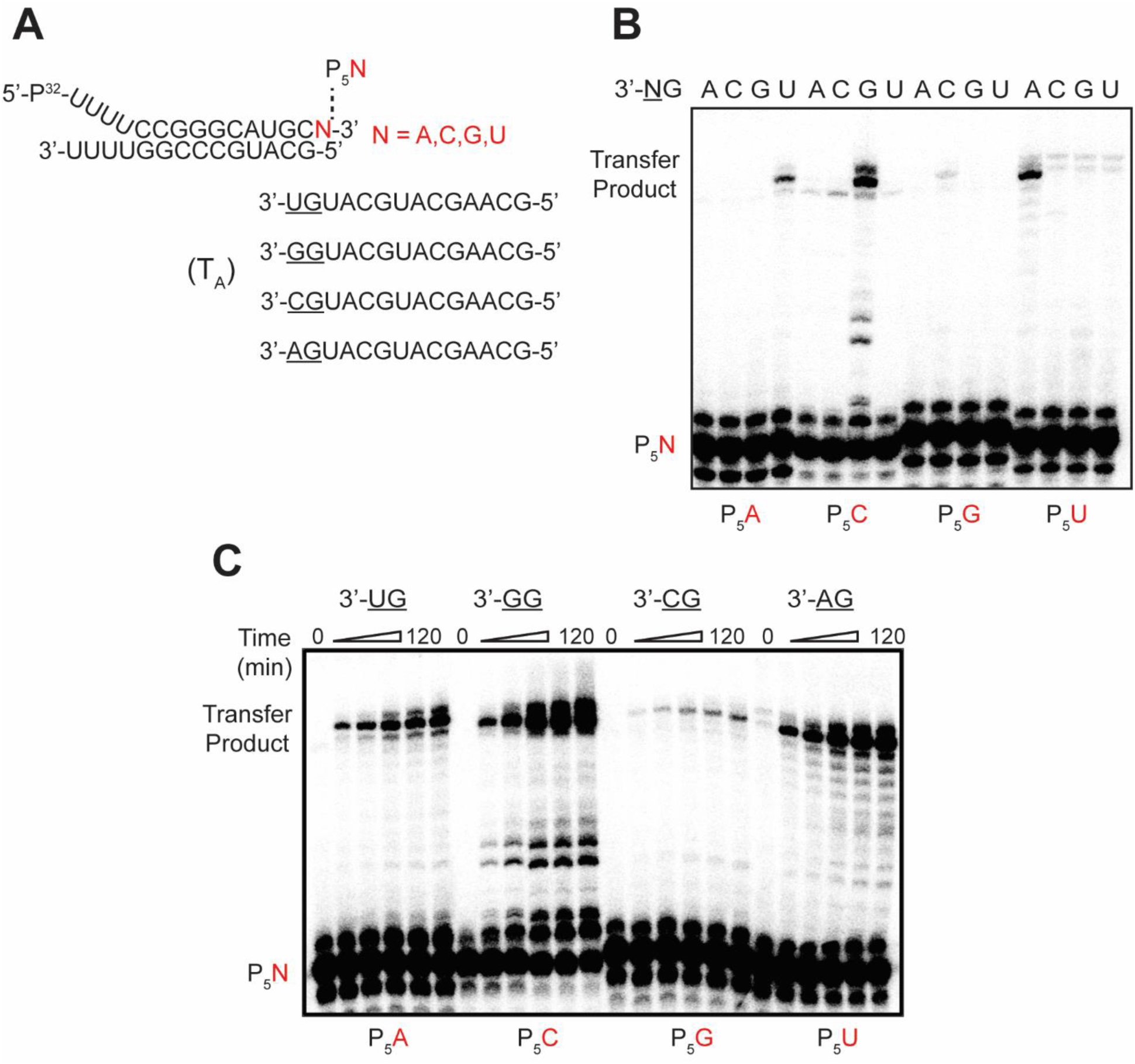
A single basepair between the intermediate and acceptor template is sufficient for template switching. P_5_N intermediates with unique 3’-terminal overhangs (N=A, C, G, and U in red) and acceptor template RNAs with unique 3’- termini (3’-UG, 3’-GG, 3’-CG, and 3’-AG) used to assess template switching by PV RdRp. (**B**) Analysis of reaction products by denaturing PAGE from template-switching reactions performed with P_5_N intermediates (P_5_A, P_5_C, P_5_G, and P_5_U) and the indicated acceptor template RNAs. The positions of the unextended primer and transfer product are indicated. Transfer product is only observed when a basepair can form between the primer and acceptor template RNA. (**C**) Comparison of the amount of transfer product RNA produced over time from template-switching reactions performed with P_5_Nintermediates and the indicated acceptor template RNAs.

**Figure 11.**
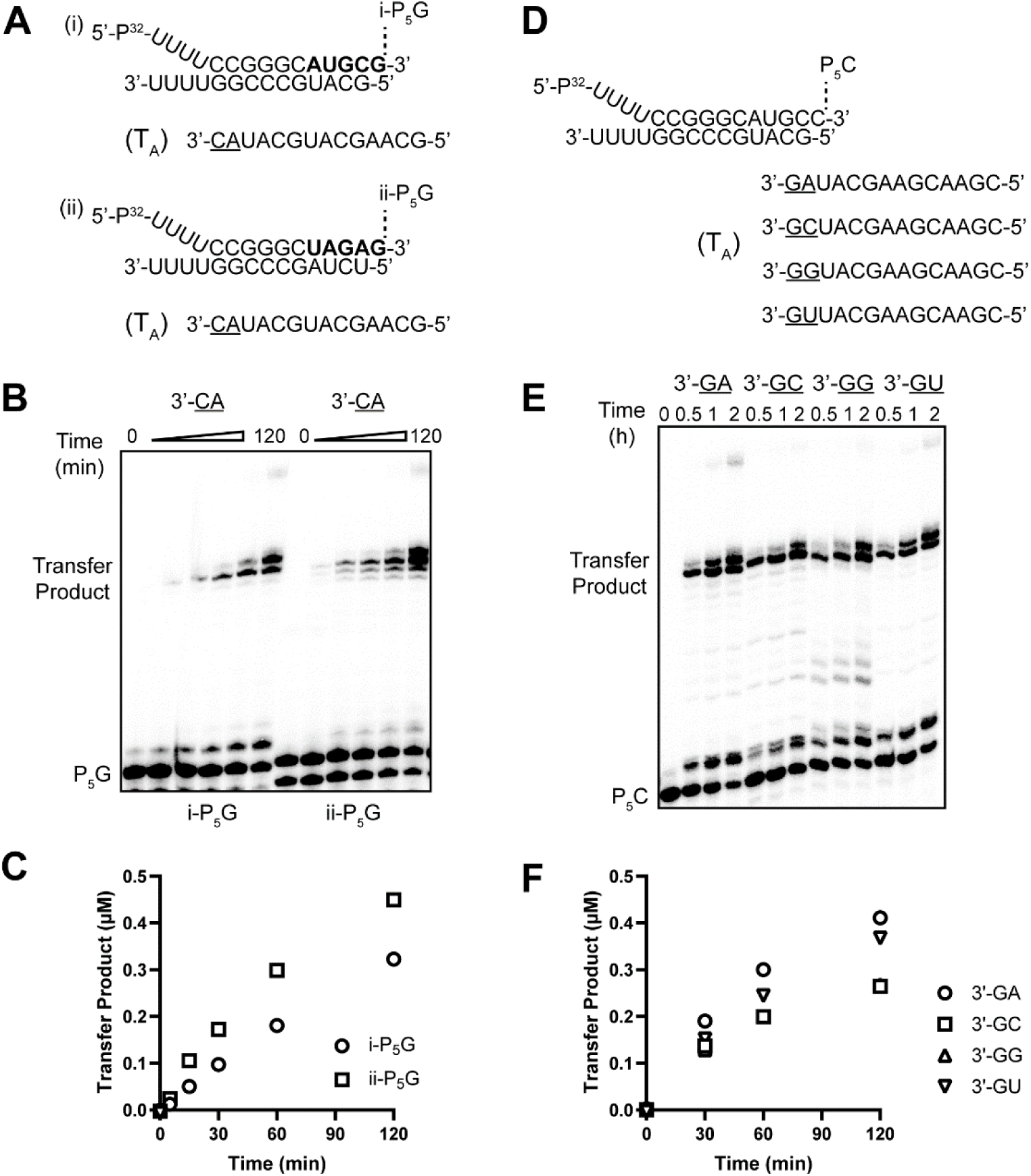
Efficiency of template switching varies depending on the sequence of the adjacent nucleotide. (**A**) P_5_G intermediates with unique 3’-terminal ends (sequence in bold is different) and acceptor template RNA used to assess template switching by PV RdRp. (**B**) Analysis of reaction products by denaturing PAGE from template-switching reactions performed with P_5_G intermediates and 3’-CA acceptor template RNA. The positions of the unextended primer and transfer product are indicated. (**C**) Quantitative analysis of the formation of transfer product as function of time from template-switching reactions performed with i-P_5_G and ii-P_5_G primed-template duplexes. (**D**) P_5_C intermediate and acceptor template RNAs with unique 3’-terminal ends (3’-GA, 3’-GC, 3’-GG, and 3’-GU ends) used to assess template switching by PV RdRp. (**E**) Analysis of reaction products by denaturing PAGE from template-switching reactions performed with P_5_C intermediate and acceptor template RNAs. The positions of the unextended primer and transfer product are indicated. (**F**) Quantitative analysis of the formation of transfer product as function of time from template-switching reactions performed with P5C intermediate and indicated 3’GN acceptor RNAs.

**Figure 12.**
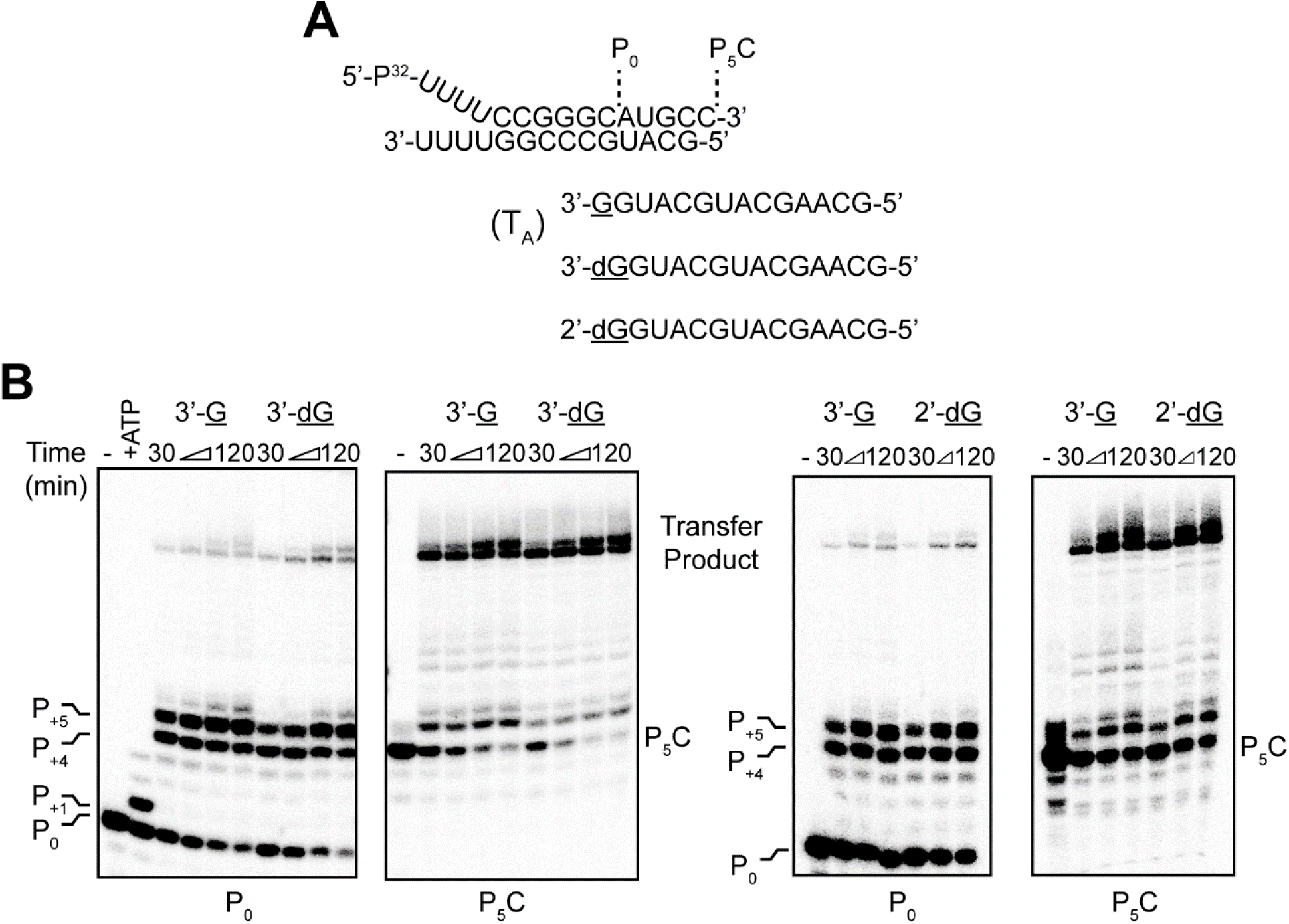
3’-deoxy and 2’-deoxy-terminated acceptor templates can serve as substrates for template switching. P_0_ primed-templates, P_5_C intermediate, and acceptor template RNAs with either a 3’-OH, 3’-deoxy, or 2’-deoxy terminated 3’-end used to assess template switching by PV RdRp. (**B**) Analysis of reaction products by denaturing PAGE from template-switching reactions performed with either P_0_ primed-templates or P_5_C intermediate and acceptor template RNAs. The positions of the unextended primer, strong-stop product (P_+4_), strong-stop, “plus-one” product (P_+5_), and transfer product are indicated.

### Observation of multiple, template-switching events

The experiments performed to this point suggested that formation of the transfer product is limited by the rate of formation and accumulation of an intermediate with a 3’-overhang complementary to the 3’-terminal nucleotide of the acceptor template. These observations suggest that the efficiency of template switching might be increased by skewing nucleotide pools to favor formation of the intermediate complementary to the acceptor template. When CTP was present at 2 mM and the other NTPs present at 20 μM, transfer product formed efficiently with the two different acceptor templates used (**Fig. 13A** and lanes labeled CTP in **Fig. 13B**). When ATP was present at 2 mM instead of CTP, template switching did not occur (lanes labeled ATP in **Fig. 13B**). We observed two template-switching events with both acceptor templates only when CTP was present at 2 mM (#1 and #2 transfer products in lanes labeled CTP in **Fig. 13B**). Interestingly, as many as five, consecutive, template-switching events occurred with the T_A2_ acceptor template (#1-#5 transfer products in lanes labeled T_A2_ and CTP in **Fig. 13B**). The difference between T_A1_ and T_A2_ is that the former terminates as follows: 3’…CG- 5’, and the latter terminates as follows: 3’…GC-5’. We synthesized a third, subtly longer acceptor template terminating with 3’…GC-5’ (T_A3_ in **Fig. 13A**). This acceptor template also supported robust template switching only in the presence of 2 mM CTP, yielding again as many as five, template-switching events (#1-#5 transfer products in lanes labeled T_A3_ and CTP in **Fig. 13B**).

**Figure 13.**
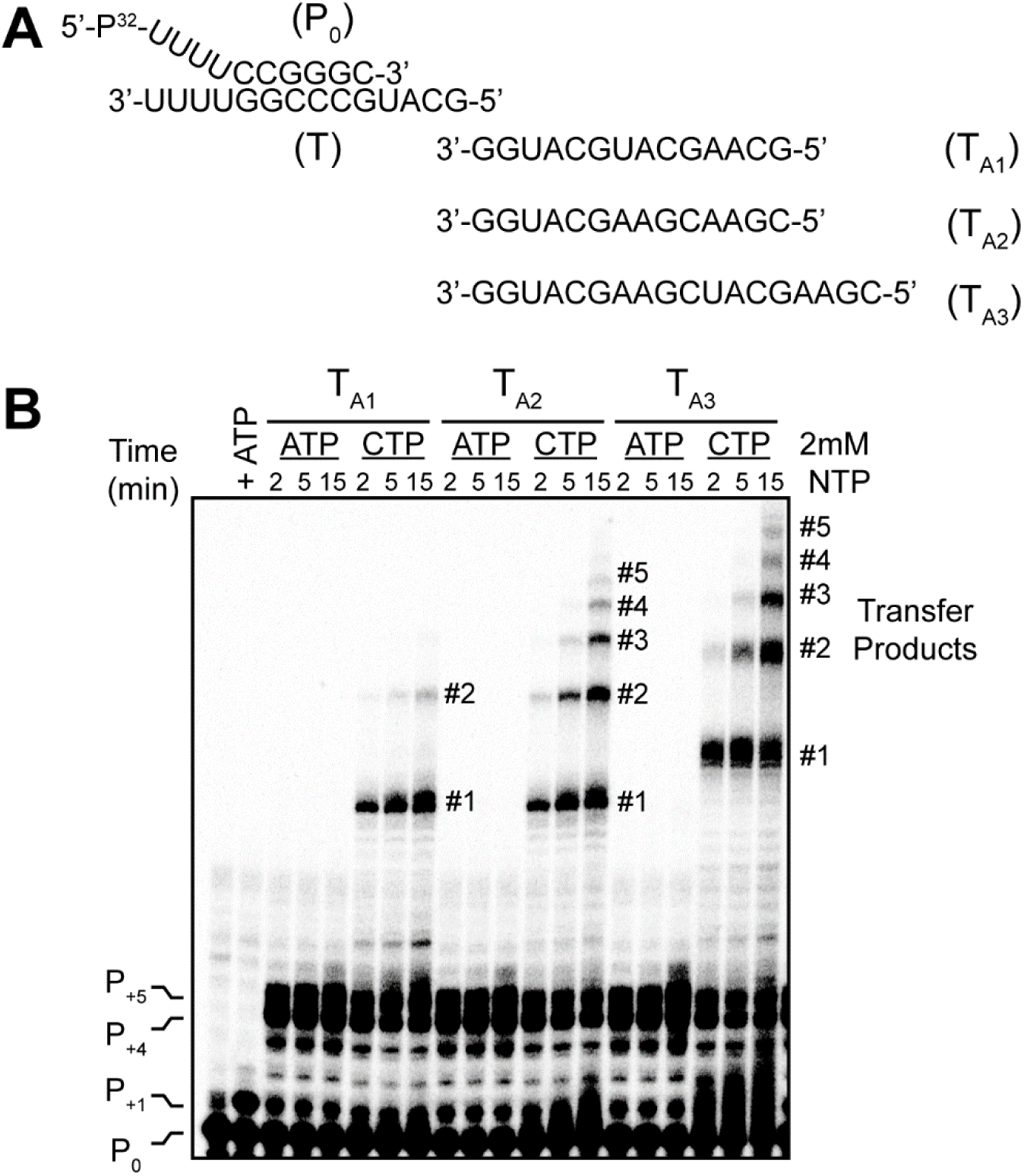
Altering the concentration of NTPs increases the frequency of template switching. (**A**) Primed-template and acceptor template RNAs (T_A1_,T_A2_, and T_A3_) used to assess template switching by PV RdRp. The sequence of the acceptor templates each have a 3’-terminal GMP residue but differ slightly in the remaining sequence. (**B**) Analysis of reaction products by denaturing PAGE from template-switching reactions performed with P_0_ primed-templates and acceptor template RNAs. The concentration of NTPs were 20 µM each nucleotide with an additional 2 mM ATP or CTP. Reactions containing 2 mM CTP promoted non-templated nucleotide addition and the ability of PV RdRp to multiple template switching events using the 3’GMP terminated acceptor template. The positions of the unextended primer, strong-stop product (P_+4_), strong-stop, “plus-one” product (P_+5_), and transfer products (#1-#5) are indicated.

### Template switching from the ends of templates resembling viral plus-and minus-strand RNAs

While contemplating the function of forced-copy-choice RNA recombination in infected cells, we considered the possibility that this process served as a last-ditch response to rescuing viable genomes in the face of extraordinary RNA damage. For example, cell-based, innate-immune mechanisms will introduce oxidants into the infected tissue, causing base modifications and perhaps even scission of the phosphodiester backbone. If this is the case, then the ability of the polymerase to distinguish between a natural 5’-end and a damaged end might be advantageous. The 5’- end of PV plus-strand RNA is a VPg-pUpU dinucleotide that is formed by using the hairpin present in 2C-coding sequence as template (**Fig. 14A**). The 5’-end of minus-strand RNA is a poly(U) stretch templated by the poly(A) tail of plus-strand RNA (**Fig. 14A**). We constructed primed-templates with 5’-ends that mimic these natural sequences and their complementary sequences (**Fig. 14B**). The design of the experiment is indicated in **Fig. 14C**. Running PV RdRp into a uridine dinucleotide or an adenine dinucleotide led to robust template switching (T-U_2_ and T-A_2_ in **Fig. 14D**). While a stretch of uridine residues presented no impediment to template switching (T-U_10_ in **Fig. 14E**), a stretch of adenine residues diminished the efficiency of formation of an elongation-competent polymerase complex as well as template switching (T-A_10_ in **Fig. 14E**). Whether or not there is something special about the presence of a poly(A) tail remains unclear and future studies will need to be designed to address this question more deeply than we can do here.

**Figure 14.**
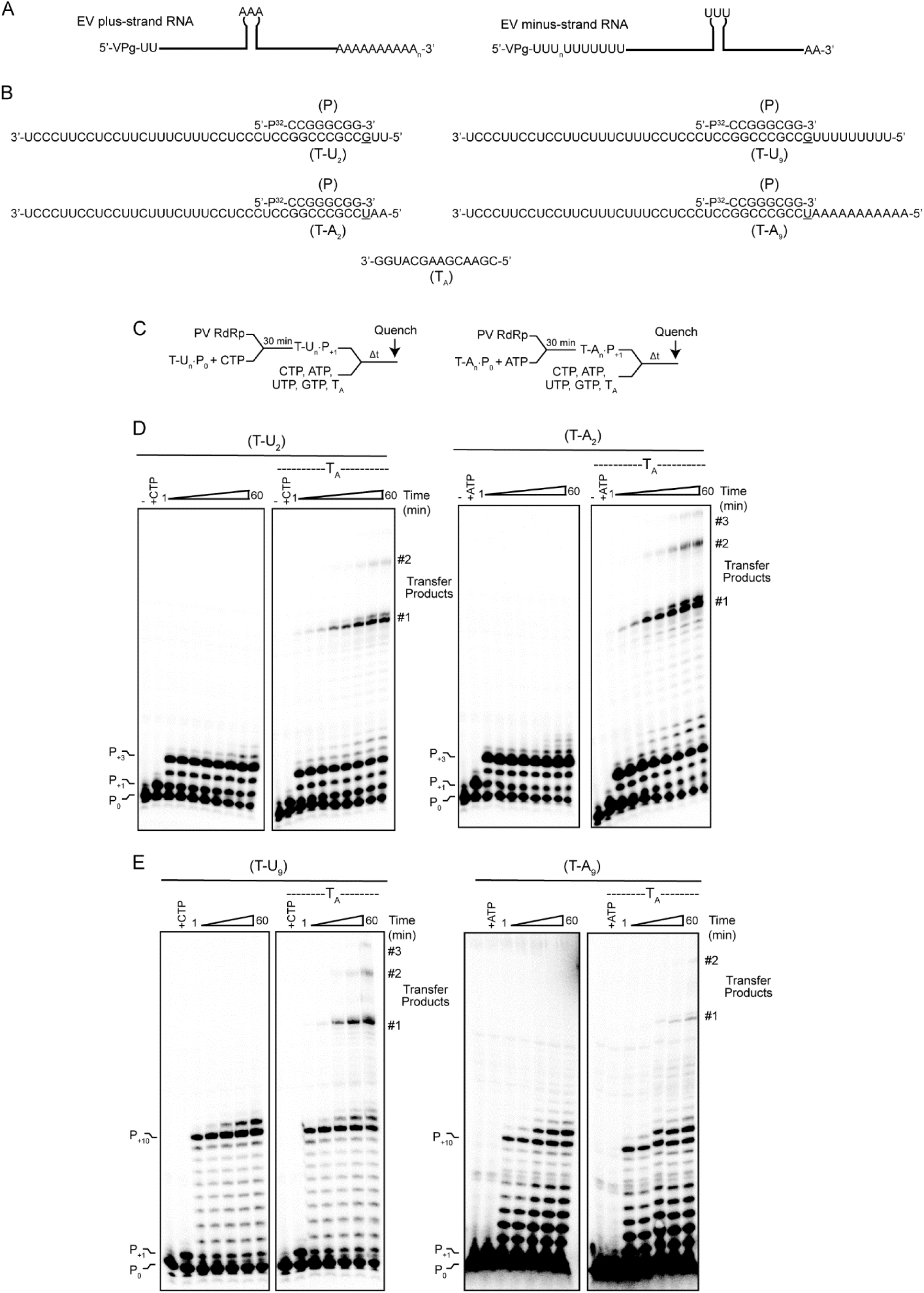
Impact of repeated nucleotides on template switching by PV RdRp. (**A**) Schematic of the enterovirus plus-strand and minus-strand RNAs. There are two uridylate residues covalently linked to VPg on the 5’-end of plus-strand and a 3’-poly-A tail of ill-defined length. There is an internal cis-replication element (CRE, oriI), that contains three adenylate residues on the apical stem-loop that serves as template to prime production of VPg-pUpU that is then transferred to the 3’-end of plus-strand RNA to produce minus-strand RNA. The 5’-end of minus-strand RNA contains VPg-pUpU and a poly-U stretch of ill-defined length; the 3’-end has two adenylates. (**B,C**) Primer and templates used in this study. The primer (P) is an 8-nt RNA and templates (T-U_2_, T-A_2_, T-U_9_, and T-A_9_) are 31-nt and 38-nt RNAs; sequences are shown. The annealed primed-template forms an 8-bp duplex with a 3-nt or 10-nt 5’-template overhang. The first templating base is underlined. The RNA primer was labeled on the 5’-end with ^32^P. (**D**) Schematic of assay. Primer extension was initiated by adding PV RdRp with primed-template in the presence of the first nucleotide to be incorporated, CTP or ATP for 30 min. After formation of P_+1_, the remaining NTPs (20 µM) and CTP (2 mM) were added in both the absence and presence of acceptor template for various amounts of time and then reactions were quenched. (**D,E**) Analysis of reaction products by denaturing PAGE using P-T-U_2_, P-T-A_2_ (panel C) and P-T-U_9_, P-T-A (panel D). The positions of the unextended primer P_0_, extended primers P_+1_, strong-stop products (P_+3_ and P_+10_), and transfer product are indicated.

### Beyond the PV RdRp

It is very clear that all picornaviruses interrogated for their ability to perform copy-choice RNA recombination in cells do so ^37^. To the best of our knowledge, this study reports the first observation of forced-copy-choice RNA recombination by a viral RdRp. While far beyond the scope of this first study to investigate the details of this mechanism of recombination for multiple picornaviruses, to maximize impact of this study, we evaluated the ability of other picornavirus polymerases to catalyze template switching by initiating from the standard primed-template (**Fig. 15**) or from the P_5_C intermediate (**Fig. 16**). The additional picornavirus RdRps evaluated were: Coxsackievirus B3 (CV-B3), enterovirus-D68 (EV-D68), human rhinovirus-C15 (RV-C15), RV-A16, and foot-and-mouth-disease virus (FMDV). There was clear evidence for template switching for all RdRps tested (**Figs. 15** and **16**). RV-A16 RdRp was by far the best at template switching (see RV-A16 in **Figs. 15** and **16**). The RdRps from CVB3, EV-D68 and RV-C15 were all efficient at primer extension (see P_+4_ in **Fig. 15**). However, production of the transfer product by these enzymes, even when starting from the P_5_C intermediate, was weak. Sorting out the basis for these differences between the various RdRps will require a study devoted specifically to this question.

**Figure 15.**
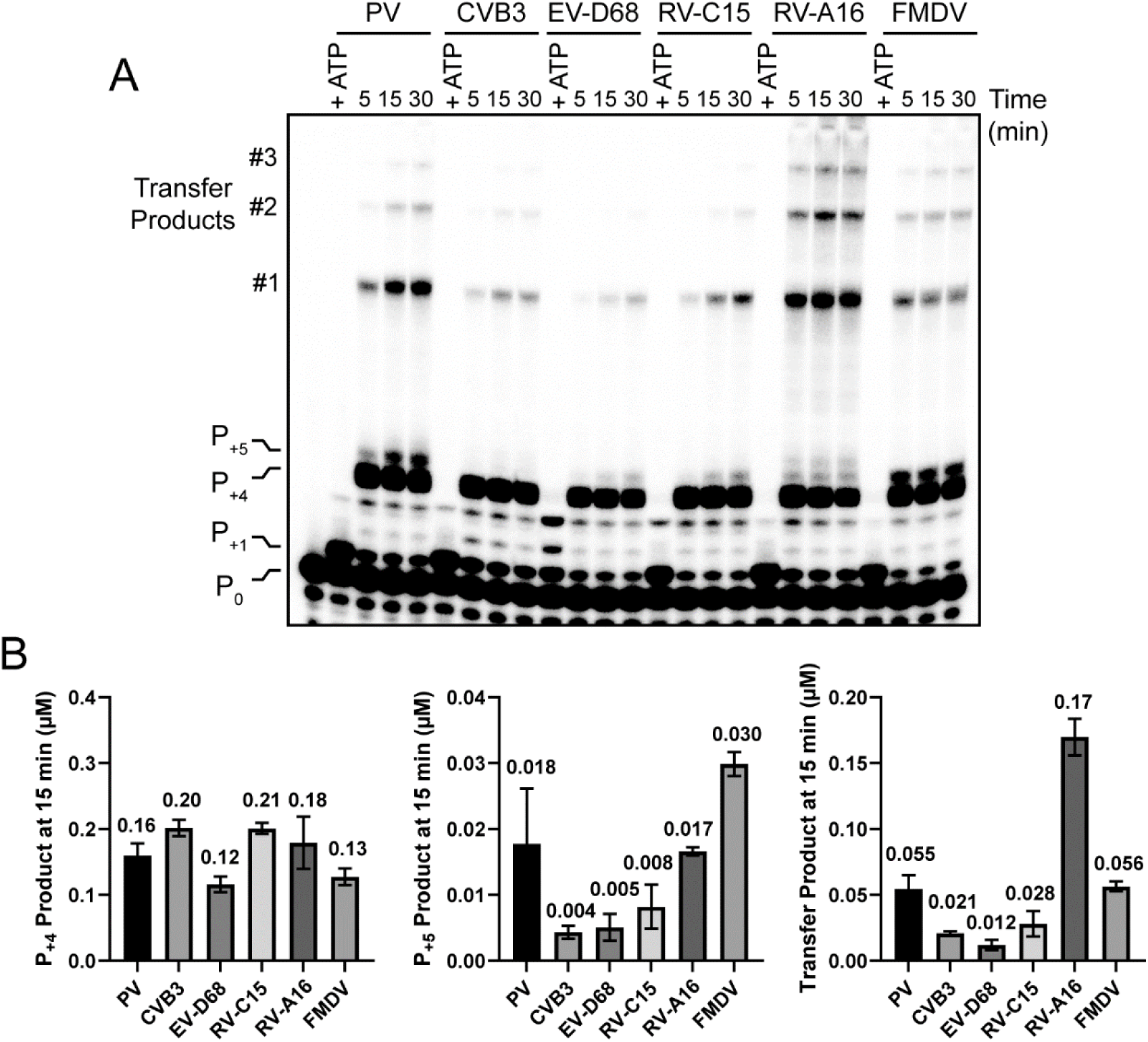
Template switching by picornaviral RdRps. (**A**) Analysis of reaction products by denaturing PAGE from template-switching reactions performed with P_0_ primed-template, acceptor template RNA, and the indicated RdRp. The concentration of NTPs were 20 µM each with an additional 2 mM CTP. Reactions promoted non-templated nucleotide addition and the ability of the RdRp to perform multiple template switching events using the 3’GMP terminated acceptor template. The positions of the unextended primer, plus-one product (P_+1_), strong-stop product (P_+4_), strong-stop, “plus-one” product (P_+5_), and transfer products (#1-#3) are indicated. (**B**) Quantitative analysis of the formation of P_+4_, P_+5_ and transfer product at 15 min from template-switching reactions performed with the indicated RdRps.

**Figure 16.**
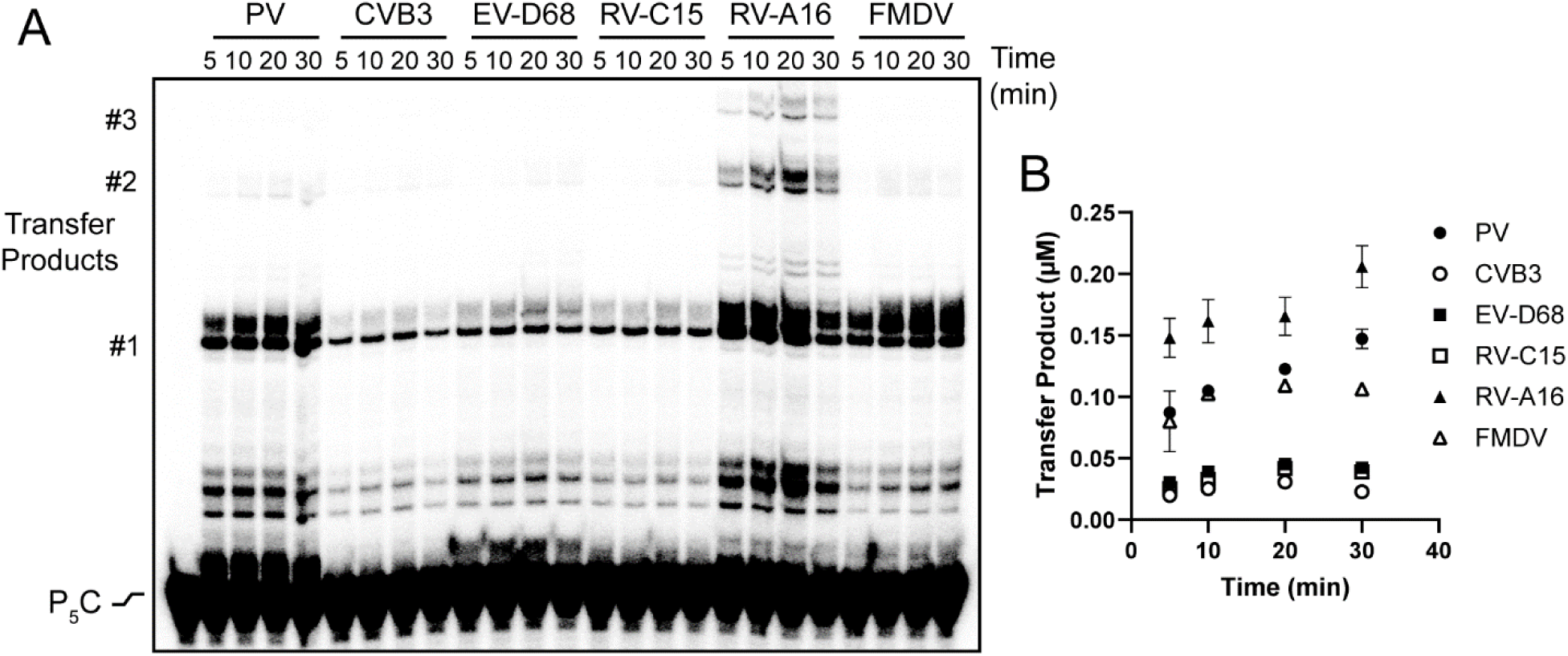
Utilization of the P_5_C intermediate by picornaviral RdRps. (**A**) Analysis of reaction products by denaturing PAGE from template-switching reactions performed with P_5_C intermediate, acceptor template RNA, and the indicated picornavirus RdRp. The concentration of NTPs were 500 µM each. Reactions promoted the RdRps to template switch. The positions of the P_5_C intermediate and transfer products (#1-#3) are indicated. (**B**) Quantitative analysis of the formation transfer products from template-switching reactions performed with the indicated RdRps.

## Discussion

Why do positive-strand RNA viruses and retroviruses recombine? Is recombination an evolved trait to maintain viral fitness of the viral population or a “mechanistic by-product” of the biochemical and biophysical properties of the viral polymerase required for genome replication^3^? The missing piece to this puzzle is the absence of a clear mechanistic description of viral RNA recombination. This circumstance is now changing^9^. Recent studies of copy-choice RNA recombination revealed an unexpected mechanism of template switching^9^. In response to nucleotide misincorporation, some fraction of elongation complexes “backtrack,” releasing the 3’-end of nascent RNA from the template. The 3’-end is then extruded through the nucleotide-entry channel into the solution, where hybridization of a complementary template can occur to complete the template-switching process^9^. The mechanism of forced-copy-choice RNA recombination reported here is equally unexpected. The use of a blunt-ended duplex with a non-templated nucleotide added to the 3’-end of one strand to form the intermediate required for template switching (**Fig. 17**). A single basepair is sufficient for this template-switching event to occur (**Fig. 17**). In both types of RNA recombination, the properties of the polymerase required for recombination are in many ways independent of the requirements for elongation and are triggered by nucleotide misincorporation. We would like to suggest here that viral RNA recombination may be an evolved trait.

**Figure 17.**
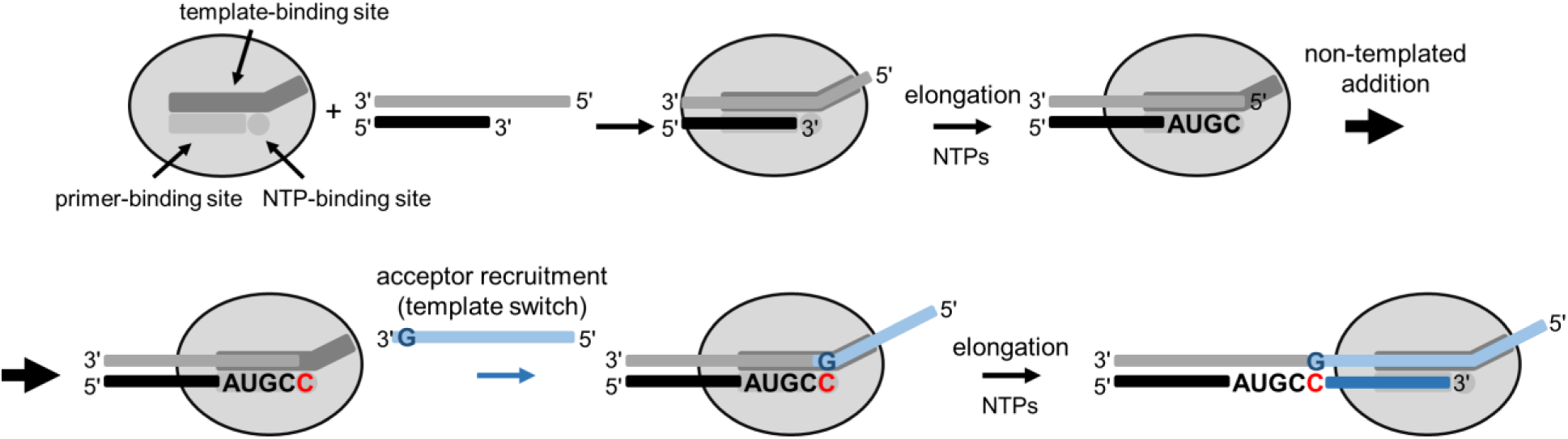
Model for enteroviral polymerase-catalyzed forced-copy-choice recombination. Mechanism for recombination as a method of rescuing viral RNA genomes following partial degradation or from abortive synthesis. Using poliovirus RdRp as a model, we have shown that the polymerase extends the nascent RNA strand to the end of its template, and then proceeds to add non-templated nucleotides. From this process a minimal 1 nt-long recombination junction is formed between the 3’-end of the nascent RNA and the 3’-end of the secondary acceptor template where elongation continues.

The very first paper published from our laboratory showed that PV RdRp was able to use template switching to make products greater than unit length when homopolymeric primed templates were used^30^. Inspired by the mechanistic studies of Peliska and Benkovic on template switching by HIV RT^17^, we added an acceptor template complementary to the 3’-end of nascent RNA produced from a self-complementary (symmetrical) primed template referred to as sym/sub^10, 34^. We demonstrated an acceptor template-dependent RNA product, formation of which was compromised by randomizing the acceptor template sequence^10^. However, efforts to sequence the recombinant products were complicated by the symmetrical nature of the starting primed template^10^. The sequencing data that we were able to obtain suggested a forced-copy-choice mechanism instead of a copy-choice mechanism^10^.

Forced-copy-choice recombination is a concept that originates from a description of retroviral reverse transcription (**Fig. 1**) ^18, 19, 21^. Initiation of genome replication is primed by a tRNA bound to the retroviral genome a few hundred nucleotides or so from the 5’-end. Initiation from the tRNA is forced to terminate when the 5’-end is reached. To continue synthesis the strong-stop product must be transferred to the 3’-end of the retroviral genome. To our knowledge, this type of recombination has never been invoked as a mechanism used by positive-strand RNA viruses. However, initiation of genome replication by enteroviruses requires a priming event that occurs at an internal position of the genome instead of the 3’-end^27^.

We established and validated a primed-template system that could be used to determine the type of RNA recombination that was occurring and the ability to elucidate its mechanism (**Figs. 1**-**5**). Non-templated addition of a single nucleotide to the 3’-end of nascent RNA once the polymerase reached the end of template was quite robust (e.g. P_+5_ in **Fig. 3C** and **Table 2**). A PV RdRp derivative (K359R) defective for non-templated addition was also defective for template switching in this assay (**Fig. 7**). This was the first indication that the P_+5_ product with the non-templated addition might be an obligatory intermediate for template switching by a forced-copy-choice mechanism. In fact, neither the blunt end nor a two nucleotides-extended product supported template switching, but the P_+5_ intermediate worked quite well for both WT RdRp (**Figs. 6** and **8**) and K359R RdRp (**Fig. 7**). A single Watson-Crick basepair was sufficient for the RdRp to match the intermediate with an appropriate acceptor template and extend the intermediate to the end of the acceptor template (**Figs. 9** and **10**) and beyond (**Fig. 13**). As is common when nucleic acid is a component of a reaction, the context in which the 3’-terminal nucleotide is presented, its nearest neighbors, modulate the efficiency of template switching (**Figs. 11** and **12**). Formation of the basepair mattered most; the nature of the sugar configuration mattered least (**Fig. 12**). Finally, all picornavirus RdRps tested were able to catalyze template switching by a forced-copy-choice mechanism whether the reaction was launched from a primed template (**Fig. 15**) or the P_+5_ intermediate (**Fig. 16**).

The mechanism of forced-copy-choice RNA recombination catalyzed by PV and other enterovirus RdRps reported here (**Fig. 17**) is essentially identical to the forced-copy-choice DNA recombination catalyzed by the reverse transcriptase encoded by a group II intron^25, 38^. Non-templated addition to a blunt-ended elongation product was essential for efficient template switching, and formation of a single basepair was sufficient for efficient template switching^25^. Different RTs are known to do this reaction, but the nature of the nucleotide incorporated and the number of nucleotides added vary^25, 38–40^. Therefore, it was quite surprising to see that both the enterovirus RdRp and group II intron RT used the exact same mechanism.

A few years ago, Lambowitz and colleagues reported the structure of the group II intron-encoded RT poised for template switching^25^. Because RT and the RdRp share substantial conservation in overall topology and conserved structural motifs and corresponding functions, we compared the RT model to a model of the PV RdRp elongation complex^41^. The PV RdRp elongation complex enabled formulation of a structure-based hypothesis for RdRp-catalyzed template switching during forced-copy-choice RNA recombination (**Fig. 18A**). The RdRp binds to the strong-stop RNA product (step 1), then adds a non-templated nucleotide (step 2), and finally recruits an acceptor template capable of baseparing with the non-templated nucleotide added (step 3). The acceptor template and nascent basepair should have the same extensive interactions experienced by the template and nascent basepair of the elongation complex (**Fig. 18B**). Importantly, this analysis revealed several residues in conserved sequence/structure motifs F (red) and G (green) that are conserved across the enterovirus RdRps that should be critical to the positioning of the acceptor template and nascent basepair for catalysis (**Fig. 18C**).

**Figure 18.**
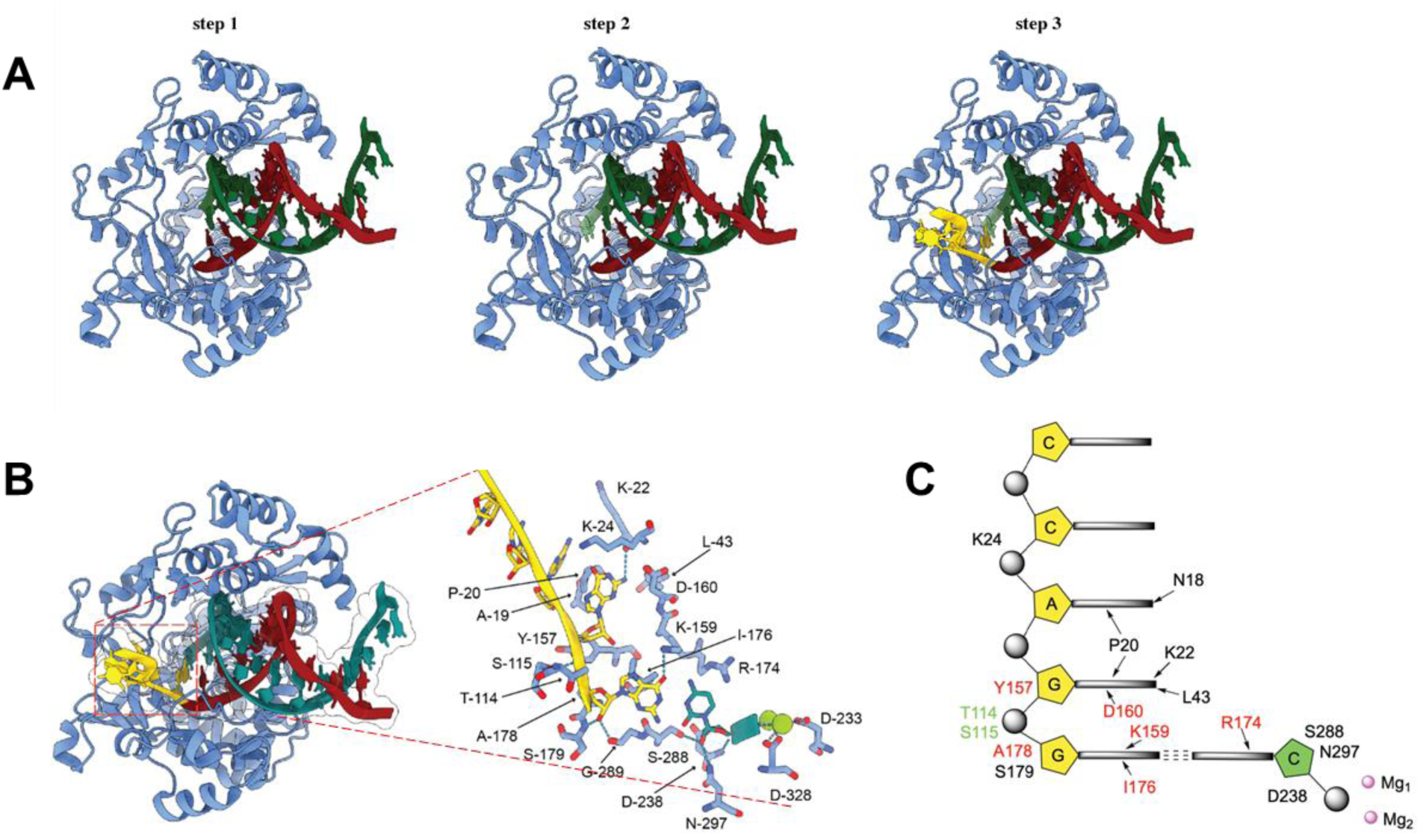
Structural model for forced-copy-choice recombination. (**A**) Using an existing structure of PV RdRp with primed template (30L7), we have produced snapshots of the proposed steps of forced-copy-choice recombination. PV RdRp extends the nascent RNA strand (green) to the end of its template (red) [step 1], and then adds a non-templated nucleotide [step 2]. Binding of the template strand (yellow) to the PV-RdRp-recombination intermediate complex leads to formation of 1-bp duplex, extension from which produces the recombinant RNA [step 3]. (**B**) Structural model for the recombination intermediate (step 3) in which PV RdRp extends the nascent RNA strand (green) 1 nt beyond the template (red) and the recombination junction is formed between the 3’-end of the nascent RNA (green) and the 3’-end of the secondary acceptor template (yellow). Boxed region showing the interactions of PV RdRp amino acid side chains with acceptor template. (**C**) Schematic of the basepairing between the P5C intermediate (green) and template (yellow). Residues stabilizing this interaction originate from conserved sequence/structural motifs F (red) and G (green)

One interesting difference between RdRp and RT-catalyzed template switching is that the RdRp appears to rarely incorporate UMP as the non-templated nucleotide but RT uses all four apparently equally^6, 17, 25, 38–40^. We were unable to observe a P_+5_ (15-nt) intermediate ending with UMP (see rows 10-12 in **Table 2**). Enteroviruses and other picornaviruses produce their proteins as a single polyprotein that is cleaved post-translationally. Inadvertent incorporation of a stop codon during recombination would be lethal to the virus. One possibility is that the inability to introduce UMP by non-templated addition may have evolved to preclude introduction of a stop codon, all of which begin with a uridine. RdRps and RTs are thought to have evolved from a common ancestor^42^. For RdRps to be incapable of adding UMP and for RTs to be capable of adding UMP must have happened after the split into the two lineages. We know that the problem is UMP addition and not utilization because a P_5_U intermediate is used by PV RdRp (**Fig. 10**). Moreover, evidence for UMP misincorporation from an internal position of template was detectable even at a frequency of 0.00009 (see row 6 in **Table 2**). This observation supports the notion that RdRp-catalyzed forced-copy-choice RNA recombination has evolved and perhaps continues to evolve independent of other known functions of the polymerase.

Another distinguishing feature of the specificity of PV RdRp for non-templated addition of nucleotides is the preference for CMP incorporation over other nucleotides (compare lines 10-12 in **Table 2**). Many RdRps have been reported to exhibit terminal ATP adenylyltransferase activity. We have assumed that this reflected the polymerase “A-rule,” which states that when in doubt, for example in the absence of a template or in the presence of a lesion in the template, polymerases incorporate AMP^43^. This rule also extends to blunt-ended duplex substrates^39^. Indeed, a commercial strategy for cloning PCR products relies on non-templated addition of AMP by Taq polymerase^44^. While the RT encoded by a group II intron only adds a single, non-templated nucleotide of any type^25^, the RT from Moloney murine leukemia virus adds three cytidine residues^40^. This observation further supports the notion that mechanisms of recombination evolved independently of the mechanism of elongation. Our data with the picornavirus RdRps support a similar conclusion. Although all RdRps tested produced equivalent amounts of strong-stop product (P_+4_ in **Fig. 15**), the efficiency of non-templated addition of nucleotides and template switching varied widely (P_+5_ and Transfer Products in **Fig. 15**). Explanation of the differences observed between RdRps will require future studies. Do all of these RdRps follow the C-rule? When the P_+5_ intermediate does not accumulate, does this mean that the addition of the non-templated nucleotide is inefficient or does this mean that recruitment of the acceptor template is more efficient?

Forced-copy-choice recombination is thought to have evolved as an RNA repair mechanism that will assemble full-length genomes from fragments^45^. The evidence for this with HIV is substantial^19, 20^. For such a mechanism to work, more than one copy of the genome is required, minimally a donor template and an acceptor template. Of course, each HIV virion contains two copies of the genome. Our studies of the RdRp show quite convincingly that the forced-copy-choice mechanism permits assembly of RNA fragments (**Figs. 13** and **15**). Now, we also know that enteroviruses spread by a non-lytic mechanism with many virions contained in vesicular carriers of different origins. Multiplicities of infection much greater than one as afforded by non-lytic spread makes an RNA repair mechanism feasible. More work in this area will be needed. For PV, we have mutants capable of only copy-choice or forced-copy-choice recombination that may be useful for exploring the role of RNA recombination in virus biology^16^.

If forced-copy-choice recombination contributes to RNA repair, then it might be advantageous to distinguish unnatural genome ends created by nucleolytic cleavage from natural genome ends (**Fig. 14A**). With the exception of an oligo(rA)-terminated RNA, all natural genome ends appeared competent for recombination (**Fig. 14**). It is unclear if the result with the oligo(rA)-terminated RNA reflects the inability to extend the corresponding duplex with this sequence. Multiple preparations yielded the same result (data not shown). More work will be required to clarify this matter.

The capacity for the enterovirus RdRp to use a one-nucleotide-extended, double-stranded nucleic acid as a primer for extension of a template and assembly of multiple fragments suggests a few applications for this reaction. First, introduction of adapters, fluorophores, peptides, anything that can be attached to an oligo can be covalently linked to the 5’-end of product RNA. We are particularly intrigued by the ability to use the forced-copy-choice mechanism of the RdRp to prepare samples for analysis by nanopore technologies. Typically, evaluation of a ssRNA by nanopore begins with the addition of adapters and use of RT to convert the ssRNA to an RNA-DNA hybrid, ending with ligase treatment to remove any nicks. The need to use RT to make a double-stranded product relates to removal of any secondary structure in the RNA that would create unique signatures when detected by the nanopore instrument. Of course, RT is not the most processive polymerase. Replacing RT with an RdRp may permit even longer templates to be evaluated and would also produce a second strand of RNA that could be analyzed, thus bolstering the reliability of interpretations made by analyzing data derived from the first strand of RNA.

## Supporting information

NGS_3A

NGS_3A_Control

## Supporting Information

Supporting information is available free of charge via the internet at http://pubs.acs.org. Supporting information contains the following: Figure S1. Comparison of assembly using primed templates with and without additional uridine residues. Table S1. The top 500 sequences from small RNA-Seq.

## Author Contributions

Conceptualization: JJA; CEC; Methodology: JJA, AM, AJ, XL, IMM and CEC; Data collection: JJA, AM, AJ, XL, and IM; Data analysis: JJA, AM, AJ, XL, IMM and CEC; Writing: JJA, AM, and CEC; Supervision: JJA and CEC; Funding acquisition: JJA and CEC;. All authors have given approval to the final version of the manuscript.

## Data Availability

All data are incorporated into the article and its online supplementary material. Constructs and data sets presented in this study are available upon request.

## Funding

This work was supported by the National Institutes of Health [R01AI045818 to C.E.C., J.J.A.] The authors declare no competing financial interest.

## Acknowledgements

We thank Dr. Andrew Woodman for his many contributions to this study. We also thank Dr. Hemant Kelkar and the UNC Bioinformatics and Analytics Research Collaborative for assistance with the small RNA-sequencing analysis pipeline.

## Abbreviations

PV: Poliovirus
RdRp: RNA-dependent RNA polymerase
RT: Reverse transcriptase
VPg: Viral protein genome-linked
CV-B3: Coxsackievirus B3
EV-D68: Enterovirus-D68
RV-C15: Human rhinovirus-C15
RV-A16: Human rhinovirus-A16
FMDV: Foot-and-mouth-disease virus
PAGE: Polyacrylamide gel electrophoresis
AMP: Adenosine monophosphate
CMP: cytidine monophosphate
GMP: guanosine monophosphate
UMP: Uridine monophosphate
EDTA: Ethylenediaminetetraacetic acid

## Notes

### Competing Interest Statement

The authors have declared no competing interest.

### Summary of Updates

update correct author names and list

