## Supplementary material for "Mechanism of forced-copy-choice RNA recombination by enteroviral RNA-dependent RNA polymerases": Fig. S1

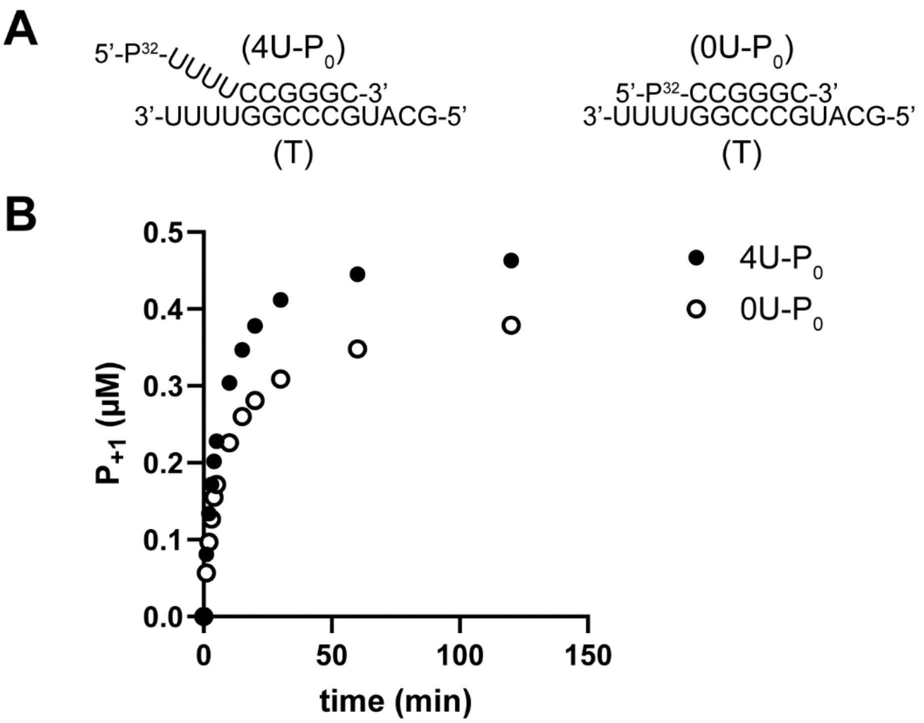


**Figure S1. Comparison of primed-templates with and without four 5'-uridine residues.** (**A**) Primed-templates with and without four 5'-uridine residues. (**B**) Quantitative analysis of the kinetics of assembly by monitoring product RNA (P+1) formation.
